# AAV gene therapy for GBA-PD and Gaucher Disease

**DOI:** 10.1101/2025.06.17.660133

**Authors:** Swathi Ayloo, Jae Cheon Ryu, Shih-Ching Chou, Lydia Blatnik, Erik Wischhof, Jie Bu, Lilu Guo, Mahmud Hossain, Dhiman Ghosh, William McCarty, Ann Byrne, Lilly Chai, Jennifer Clarke, Can Kayatekin, Bailin Zhang, S. Pablo Sardi, Martin Goulet, Bradford Elmer, Christian Mueller, Shyam Ramachandran

**Author notes:** Correspondence should be addressed to CM.

## Abstract

Mutations in *GBA1*, the gene encoding glucocerebrosidase (GCase), are the most common risk factor for Parkinson’s Disease (PD). GBA-PD patients are a genetic subpopulation of PD carrying heterozygous mutations in *GBA1*. Additionally, bi-allelic mutations in *GBA1* cause Gaucher Disease (GD), a lysosomal storage disorder. Loss of GCase activity, a lysosomal enzyme leads to the accumulation of lipid substrates, disrupting lipid homeostasis and promoting cellular toxicity. Here, we report an AAV-mediated *GBA1* replacement strategy to treat GD and GBA-PD by a one-time infusion via intravenous (GD Type 1) or intra-CSF (GBA-PD) route of administration. We engineered human GCase to be readily secretable to facilitate broad cross-correction. We developed CBE (conduritol ß-epoxide) induced lipid accumulation models to assess efficacy in mice and non-human primates (NHPs) to assess efficacy of our engineered constructs. Based on data across species, across different routes of administration, we nominated AAV.GMU01 SS3-GBA1 as our lead candidate. SS3-GBA1 is robustly secreted, cross-corrected across tissues and promotes lipid clearance. By comparing human GCase levels in AAV-treated NHP brains to healthy human donor brains, we demonstrate that AAV.GMU01 SS3-GBA1 replenishes the GCase deficit seen in GBA-PD patients, thus, restoring GCase to near-physiological levels Importantly, AAV.GMU01 SS3-GBA1 is well-tolerated with no adverse findings. Collectively, we establish a therapeutic strategy for the treatment of Gaucher Disease and GBA-PD with a single gene therapy product.

**One Sentence Summary:** A novel gene therapy strategy for GBA1-PD and Gaucher disease with an engineered payload that robustly cross-corrects enhancing therapeutic footprint

## Introduction

Glucocerebrosidase (GCase), encoded by the gene *GBA1,* is a ubiquitously expressed lysosomal enzyme that breaks down lipid substrates, glucosylceramide (GL-1) and glucosylsphingosine (Lyso-GL1) into glucose and ceramide. Loss-of-function mutations resulting in reduced GCase activity cause lipid accumulation within lysosomes, leading to lysosomal dysfunction, thus disrupting tissue homeostasis (*1*). Bi-allelic mutations in *GBA1* cause Gaucher Disease (GD), the most prevalent lysosomal storage disease. GD patients generally present with hepatomegaly and splenomegaly (enlarged liver and spleen), joint pain with weak muscles and osteoporosis with bone complications. Liver, spleen, muscles and bone marrow manifestations are largely due to “Gaucher” cells that infiltrate these tissues with massive buildup of lipids in them (*2*). The accumulation of lipids is in turn due to lack of GCase enzymatic activity. Over the past 20 years epidemiological studies in different GD patient cohorts identified a strong genetic link between mutations in the *GBA1* gene and Parkinson’s Disease (PD) (*3–7*). This genetic risk was definitively established through a large, international, multi-center study which estimated an odds ratio exceeding 5, implicating mutations in the *GBA1* gene as the most common risk factor for PD (*8*). Subsequent studies also demonstrated an earlier disease onset (*9*) and more aggressive disease progression (*10, 11*) in PD patients that were carriers of *GBA1* mutations, referred to as GBA-PD patients from here on.

PD is a progressive neurodegenerative disorder that causes motor and cognitive deficits in patients. A hallmark feature of PD is the loss of dopaminergic neurons in the substantia nigra and in most cases diagnosis happens when >50% of these neurons are already lost in the brain (*12*). Pathological accumulation of α-synuclein, as demonstrated by α-synuclein quantification assays, is a characteristic biomarker of GBA-PD patients (*13*). Brain tissue analyses of PD patients also reveal that decreased GCase activity and GBA1 protein levels (*14*) are correlated with increased α-synuclein (*15*). While the mechanistic link between lipid homeostasis and α- synuclein pathology is not fully understood, a picture is beginning to emerge. Increased lipid substrates have been shown to promote α-synuclein oligomerization and aggregation (*16–18*). Indeed, recent lipidomic analyses of plasma, cerebrospinal fluid (CSF), and brain tissue samples from GBA-PD patients have now revealed significant alterations and increases in glucosphingolipids (*19–21*). Our previous work established that augmenting GCase activity via expression of functional *GBA1* gene in the central nervous system (CNS) promotes substrate clearance and reduces both GBA-related as well non-GBA associated synucleopathies (*22, 23*). Targeted delivery of the GBA1 gene to the hippocampus increased GCase activity, reduced α- synuclein aggregation and improved cognitive deficits in both neuronopathic GD (*22*) and PD-related synucleinopathic (*23*) mouse models. These studies demonstrate proof-of-concept that restoring functional GCase promotes substrate clearance and mitigates neuronal dysfunction. Importantly, these findings highlight the potential of GCase expression to be a disease-modifying therapeutic strategy. Building on these findings, Prevail Therapeutics (subsidiary of Eli Lilly) is developing an AAV9-based gene therapy for GBA-PD patients (*24*). On the other hand, for GD patients the current standard of care involves bi-weekly infusion of recombinant GCase, called enzyme replacement therapy (ERT). Although ERTs successfully mitigate disease progression and help GD Type 1 patients with symptom management in GD Type 1, not all tissues are effectively targeted such as lung and bone marrow. Additionally, patients carry the disease burden of performing multi-hour infusions on a bi-weekly basis throughout their lives. Recently, Freeline Therapeutics (WO2019070894A1, WO2022023761A2) has administered one-time AAV-GBA1 treatment in GD Type 1 patients and has reported the safety and efficacy of their gene therapy strategy. Key aspects of our strategy that differentiate us are: use of a novel CNS tropic capsid, AAV.GMU01 (*25*) as opposed to AAV9 used by Prevail Therapeutics; and engineered human GCase that is different from the mutated version of GCase used by Freeline Therapeutics.

In this study, we report the development of a novel AAV-based gene therapy strategy applicable to both GBA-PD and to GD. Our approach is distinct from those employed by both Prevail Therapeutics and Freeline (now Spur Therapeutics), as it combines a novel capsid, AAV.GMU01 (*25*) (WO2021/102234), expressing an engineered, secretable human GCase (patent application WO2025106878A1) to facilitate cross-correction across multiple tissues, thereby establishing a broader therapeutic footprint. The AAV.GMU01 capsid demonstrates broad biodistribution in the CNS when administered via the intra-cerebrospinal fluid (intra-CSF) route in non-human primates (NHPs) while also efficiently targeting peripheral tissues when delivered intravenously (IV). These characteristics make AAV.GMU01 well-suited for gene delivery in GBA-PD and GD via intra-CSF and IV administration, respectively. To further validate our strategy and demonstrate therapeutic efficacy, we developed an efficacy model both in mice and NHPs to induce lipid substrate accumulation, to mimic disease pathology seen in patients. Through a comprehensive series of pharmacological and efficacy studies in both murine and large animal models, we demonstrate that this innovative gene therapy approach has the potential to provide robust and durable therapeutic benefits for both GD and GBA-PD and could extend to idiopathic PD. These results set the stage for the advancement of this strategy as a transformative treatment for GBA-PD and GD.

## Results

### Engineering of huGBA1 protein to be readily secretable

PD is a neurodegenerative disease affecting multiple brain regions including the frontal cortex, amygdala, striatum, thalamus, midbrain substantia nigra, cerebellum and brainstem (*26*). Therefore, an effective therapy must target the brain broadly across multiple regions. To achieve this, we engineered human GBA1 protein (huGBA1, NP_000148.2) to be readily secretable to enable cross-correction, a process where secreted extracellular proteins are taken up by neighboring cells, as observed with several lysosomal enzymes. Since AAV-based gene therapy is limited by the extent of AAV transduction, engineering a secretable payload offers an advantage by expanding the therapeutic footprint beyond transduced cells, maintaining low clinical doses.

To enhance GBA1 secretion, we replaced the endogenous signal sequence at the N-terminus of human GCase protein with signal sequences from highly secreted proteins, without altering the mature GCase protein amino-acid sequence. This strategy ensures that, once cleaved by the signal peptidase complex in the endoplasmic reticulum, the mature GBA1 protein remains identical to the native human sequence (*27*). Using a combination of *in silico* tools (SignalP6.0 and DeepLoc1.0 - see Methods), we identified 4 signal sequences (SS1 through SS4) with high probabilities of secretion and signal sequence cleavage (> 96%) (**Figure 1A**).

**Figure 1.**
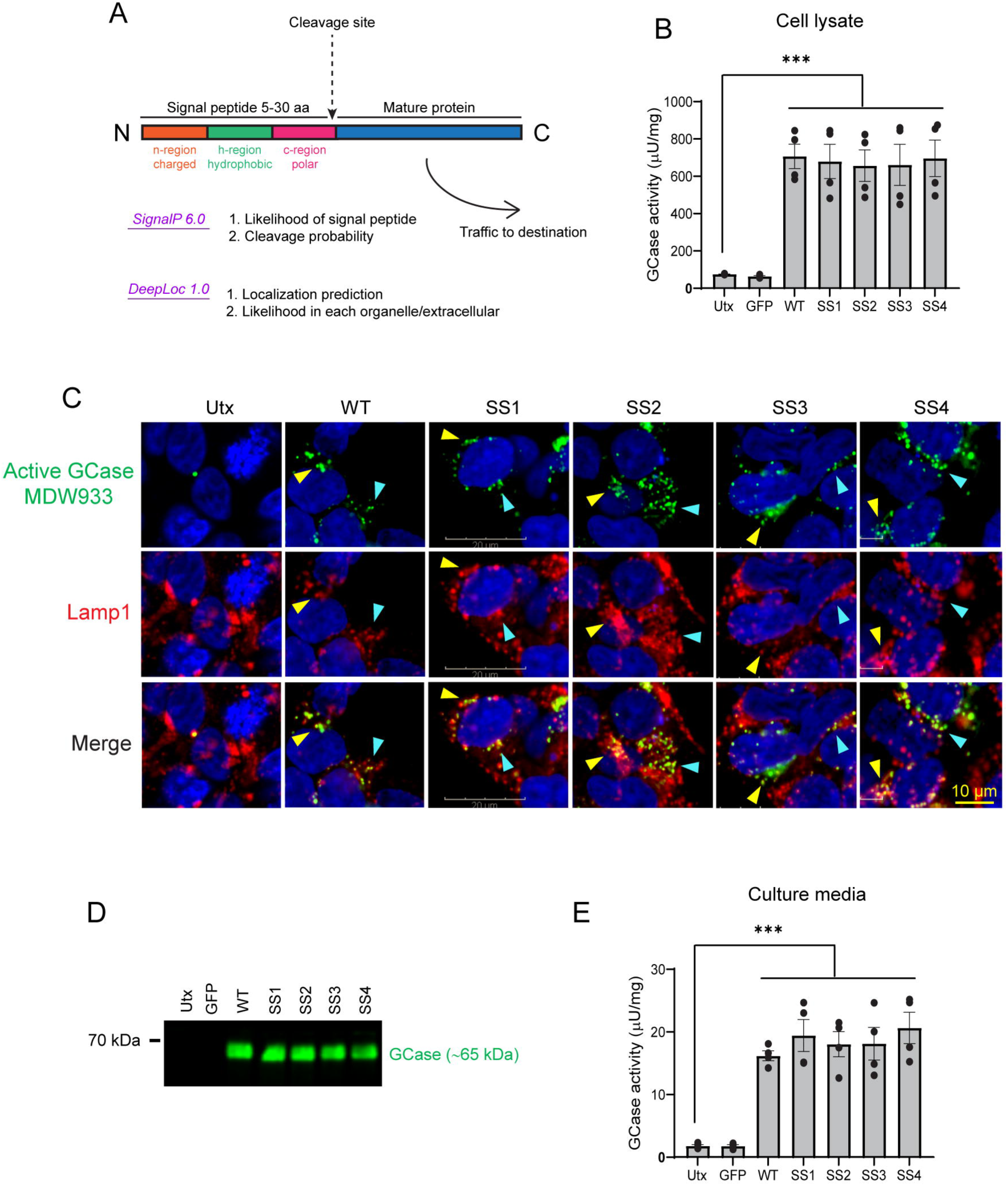
Strategy to generate engineered *GBA1* variants and *in vitro* validation of constructs. (A) A schematic depicting our strategy to generate *GBA1* variants with enhanced secretion. Endogenous signal sequence of human *GBA1* was swapped with signal sequences from highly secreted proteins. Top 4 signal sequences (SS) were narrowed down using in-silico tools that predicted robust secretion as well as high (>96%) probability of cleavage at the end of signal sequence. (B) HEK293T cells were transfected with *GBA1* plasmid containing the indicated SS variants, lysed, and GCase enzyme activity was determined in cell lysates. Mean ± SEM, n = 4 independent experiments. Untransfected versus all *GBA1* constructs as well as GFP transfected versus all *GBA1* constructs: ***p<0.001; One-way ANOVA with Turkey’s multiple comparison test. (C) Representative immunofluorescence images demonstrating co-localization of active GCase and lysosomes. HEK293T cells were transfected with *GBA1* variant constructs, incubated with MDW933 fluorescence probe to label active GCase, and immunostained with Lamp1 antibody for lysosomes. Blue and yellow arrowheads point to individual puncta showing co-localization of GCase with Lamp1. Scale bar is 10 microns. (D and E) HEK293T cells were transfected with *GBA1* variant constructs and cell culture media was collected to detect secreted GCase (D) and measure GCase enzyme activity (E). Mean ± SEM, n = 4 independent experiments. Untransfected versus all *GBA1* constructs as well as GFP transfected versus all *GBA1* constructs: ***p<0.001; One way-ANOVA with Tukey’s multiple comparisons test.

### Functional validation of engineered *GBA1* variants

To ensure that replacing N-terminal signal sequences of wildtype GCase (WT GBA1) did not alter protein function, we first evaluated enzyme activity in HEK293T cells transfected with either WT or engineered GCase variants. WT *GBA1* transfected HEK293T cells showed significantly increased GCase enzyme activity compared to untransfected cells and the enzyme activity across all GCase variants was comparable to that of WT *GBA1* with no statistically significant differences (**Figure 1B**). We then performed fluorescence imaging to examine the sub-cellular localization of the GCase variants. We used MDW933, a fluorescent probe that only binds active GCase (*28*) and observed that all variants are enzymatically active, and co-localize with lysosomes, as indicated by colocalization with Lamp1 (**Figure 1C**). Thus, engineering our *GBA1* payload to be readily secreted did not affect the enzymatic activity or the sub-cellular localization, indicating functional GBA1 protein.

We evaluated secretion both *in vitro* and *in vivo*. *In vitro,* we transfected HEK293T cells with *GBA1* variants and measured human GCase protein levels and enzymatic activity in cell culture media. Similar amount of secreted human GCase protein was detected from both WT and all the engineered GCase variants in the culture media (**Figure 1D**) and the enzymatic activity in the collected media was comparable between the WT and variant GCase constructs (**Figure 1E**). In parallel, we also executed a mouse study to assess secretion *in vivo*. 4-month-old WT mice received bilateral ICV (intra-cerebroventricular) injections of the different AAV-GBA1 variants and brain sections were analyzed after 4 weeks of AAV expression (**Figure 2A**). We performed *in situ* hybridization against the transgene (using WPRE probe) to evaluate distribution of AAV-transduced cells. To correlate AAV-transduction with protein distribution, we performed immunohistochemistry for human GBA1 protein on adjacent fixed sections. While the AAV largely transduced regions around the ventricles in the sagittal sections of all treatment groups, we observed a much larger spread of the GCase protein in SS1, SS2, SS3 compared to the WT GCase (**Figure 2B**). Higher magnification imaging of these sections revealed robust secretion of the engineered GCase variants, in this case SS3-GBA1 shown as a representative example (**Figure 2C**). Additionally, we also observed efficient uptake of the secreted protein in both neuronal and non-neuronal cell types in the brain as seen in the morphological differences of cells taking up the secreted protein (marked by green arrows).

**Figure 2.**
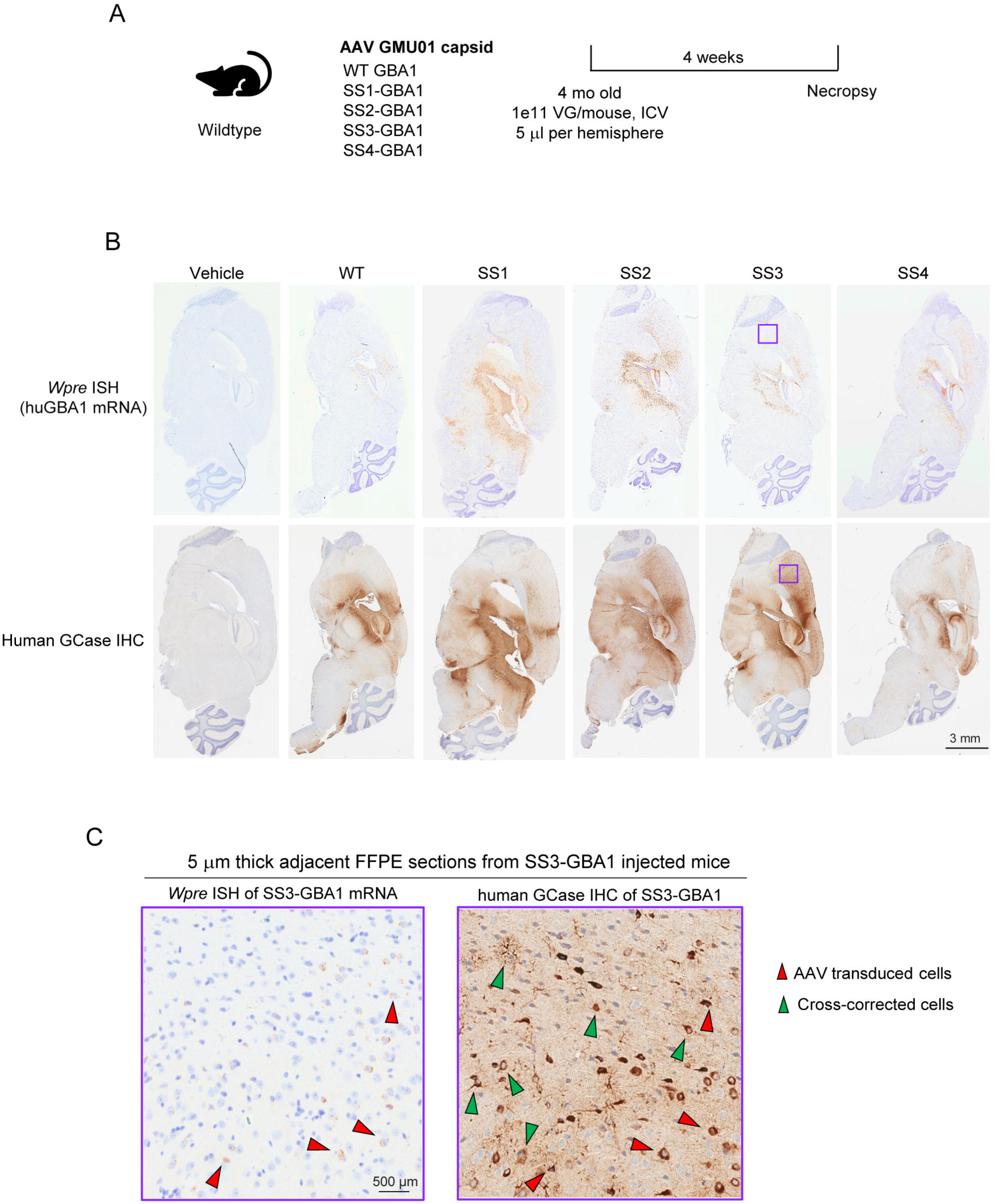
Robust secretion and cross-correction of GBA1 variants with AAV GMU01 capsid *in vivo*. (A) Study design: AAV GMU01 capsid expressing the indicated *GBA1* variants were administered by bilateral ICV to 4 month-old C57/BL6 mice at 1e11 vector genomes per animal and 5 μl per hemisphere. N=4 animals per group. The sagittal sections of brain hemisphere were analyzed 4 weeks post-injection. (B) Representative images demonstrating GCase secretion with engineered variants. Vector biodistribution is shown with *in situ* hybridization to WPRE (top panels) and GCase is shown with immunohistochemistry. (C) Representative images demonstrating cross-correction. Higher magnification images of WPRE mRNA (left) and GCase protein (right) in SS3-GBA1 injected mice brain, image corresponds to the purple box in (A). Cells positive for both WPRE mRNA and GCase are shown in red arrows (AAV-transduced cells) while mRNA negative and GCase positive cells are shown in green arrows (cross-corrected cells). Scale bar is 500 microns.

### SS3-GBA1 promotes effective lipid clearance in WT mice

To rank order the engineered *GBA1* variants by functional activity *in vivo*, we developed a mouse model of glycosphingolipid accumulation to enable evaluation of substrate reduction following treatment with our GBA1 variants. To induce lipid accumulation, we utilized CBE (conduritol ß-epoxide) which is an irreversible, brain penetrant inhibitor of GCase and has been widely used to inhibit GCase enzymatic activity across studies (*29*).

We first performed a dose response to CBE in WT mice, and measured the remaining GCase activity 24 hours post CBE administration **(Figure S1A)**. Beyond 10 mg/kg in the brain and beyond 30 mg/kg in the liver, we observed little to no change in the inhibition of GCase activity. However, when we performed lipidomics to quantify Lyso-GL1, we observed a dose-dependent increase in Lyso-GL1 accumulation in both cortex and liver, with 100 mg/kg of CBE yielding the highest accumulation of Lyso-GL1 (**Figure 3A**). These findings suggest that the GCase activity assay likely has a floor effect and Lyso-GL1 is a more sensitive and accurate read-out to query functional GCase activity at high concentrations of CBE. To further validate CBE inhibition in a more disease relevant disease model, we also investigated changes in Lyso-GL1 after 100 mg/kg CBE administration in mice harboring mutations in *GBA1* (*30–32*). In addition to WT mice, we examined two mouse lines: *Gba^D409V/D409V^, Gba^D409V/+^* which are homozygous or heterozygous for a loss-of-function D409V mutation in *GBA1*. Here we examined Lyso-GL1 in the cortex and liver at 3-, 6-, 24-, 48- and 72-hours post dosing (**Figure 3B**). With this time-course experiment, we could measure kinetics of Lyso-GL1 lipid accumulation and its subsequent clearance. Across all three genotypes, we observed highest lipid accumulation at 24 hours post CBE and then a slow reduction over the course of next 2 days (**Figure 3B**). As expected, WT mice restored lipid substrates to physiological levels the fastest, followed by the heterozygous and then the homozygous mice. Given that we observed highest lipid accumulation at 24-hour timepoint post-dosing, we used this CBE dosing regimen (100 mg/kg dose, 24 hours prior to necropsy) to evaluate the efficacy of our engineered GCase variants in promoting lipid clearance when compared to buffer-injected mice. We injected 5 different AAVs via bilateral ICV (1e11 VG per mouse, 5 ul per hemisphere) into 4-month-old WT mice, allowed AAVs to express for 4 weeks and evaluated lipid-clearance 24-hours post CBE dosing (**Figure 3C**).

**Figure 3.**
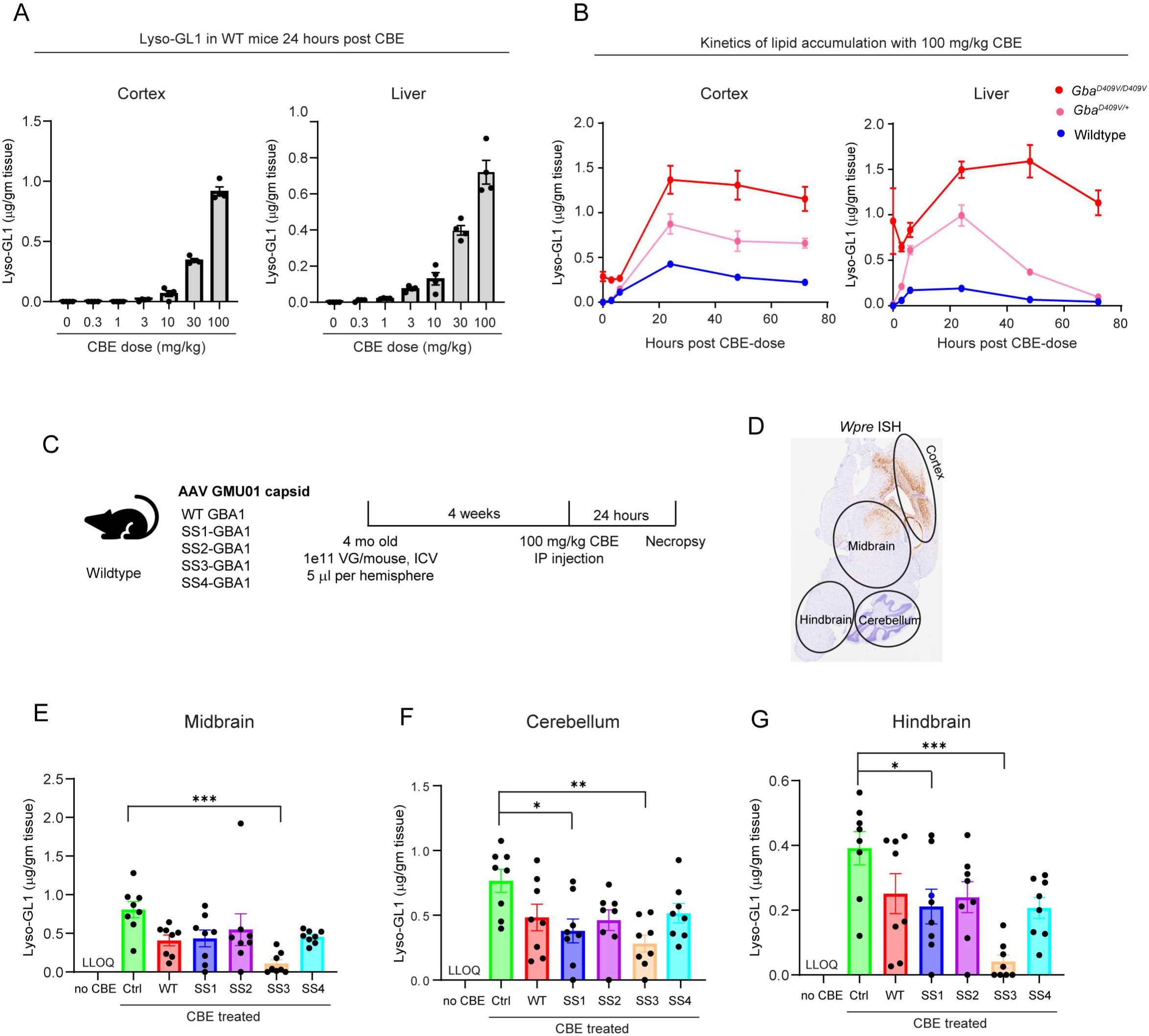
CBE-induced Lipid flux and efficient lipid substrate clearance by SS3-GBA1 in WT mice. (A) Measurement of Lyso-GL1 lipid substrate levels by LC-MS in cortex and liver of C57/BL6 mice 24 hours post CBE injection by IP injection at indicated doses. N=7 animals for CBE 0 mg/kg and N=4 per all the other groups. Data are Mean ± SEM. (B) Kinetic analysis of Lyso-GL1 lipid substrate by LC-MS in cortex and liver of 8-month-old mice from three different genotypes (*Gba^D409V/D409V^, Gba^D409V/+,^ and Gba^+/+^*). Mice with 100 mg/kg CBE IP dosing. Data are Mean ± SEM. 8 – 10 animals at each time point. (C) Study design: AAV GMU01 capsid expressing the indicated *GBA1* variants were administered by bilateral ICV to 4-month-old C57/BL6 mice at 1e11 vector genomes per animal and 5μl per hemisphere. N=8 animals per group. CBE (conduritol β-epoxide) was administered via IP injection at 100 mg/kg 24 hours prior to necropsy. (D) Representative image of *in situ* hybridization to WPRE mRNA from the analyzed sagittal sections. Cortex is proximal to site of injection and cerebellum is distal to site of injection. (E-G) Measurement of Lyso-GL1 lipid substrate levels by LC-MS in midbrain (E), cerebellum (F), and hindbrain (G). No CBE group in the graphs shown is control mice that did not receive any AAV vector or CBE injection. 8 mice per group. ***p<0.001, **p<0.01, *p<0.05; One way- ANOVA with Tukey’s multiple comparisons test with all groups compared to CBE-treated vehicle injected group.

For cross-correction to work efficiently, *GBA1* variants should be robustly secreted, taken up by non-transduced cells and maintain cellular function in these cells. We determined efficacy of cross-correction by examining lipid clearance in the distal brain regions, as a functional read-out of GBA1 protein. To assess cross-correction, we micro-dissected the brain into smaller regions of cortex, (region proximal to site of injection), midbrain, and cerebellum, hindbrain (regions distal to site of injection) (**Figure 3D**) to investigate lipid clearance in proximal and distal brain regions, as a functional read-out of GBA1 protein. We first determined vector exposure in these regions, and consistent with the route of administration of the vector, we observed higher vector transduction in cortex and midbrain and an order of magnitude lower vector genomes (VGs) in distal brain regions of cerebellum and hindbrain **(Figure S1B)**. Importantly, VGs across all AAV-injected groups were comparable indicating equivalent vector exposure across all our *GBA1* variants. As expected, compared to naïve WT mice, CBE-injected mice had a significant increase in Lyso-GL1 across all brain regions. Supporting a role for *GBA1* in clearing Lyso-GL1, all mice injected with either WT GBA1 or variants had a trend of reduced Lyso-GL1 compared to control mice injected with vehicle. However, not all variants demonstrated superiority over WT GBA1; only SS3-GBA1 had a significant effect in reducing Lyso-GL1 across all brain regions (**Figure 3E-G**). SS3-GBA1 was extremely potent in clearing substrates: in some mice, CBE-induced increased Lyso-GL1 returned to undetectable levels or below the lower limit of quantification (LLOQ), as shown in datapoints plotted as 0 in the hindbrain and midbrain tissues.

SS3-GBA1 not only had a robust effect in clearance of Lyso-GL1 lipids, it also promoted clearance of GL1 lipids **(Figure S1C-E)**.

Collectively, our data reveal that our payload strategy increases huGBA1 protein secretion without altering GCase activity or its localization to lysosomes. Our strategy enabled robust cross-correction with the secreted protein being taken up by multiple cell types as seen in our histological analyses. Our functional data further demonstrates that all variants including WT GBA1 reduce Lyso-GL1. However, SS3-GBA1 outperformed both WT GBA1 and all the other variants. SS3-GBA1 was the only variant that reduced both Lyso-GL1 and GL1, and the only variant that performed consistently well across all brain regions surveyed. Based on these data, we nominated SS3-GBA1 as our lead candidate and evaluated its efficacy and safety in subsequent NHPs studies.

### Development of CBE-induced lipid flux model in NHPs

The CBE lipid flux model developed in rodents provided the foundation for us to establish a similar model in NHPs which could then be used to run efficacy studies in a more translationally relevant model. Using allometric scaling of drug doses across species, we injected 3, 10 or 30 mg/kg of CBE into cynomolgous monkeys and collected brain, liver and plasma for lipid analyses, 24 hours post CBE-dosing (**Figure 4A**). Lyso-GL1 went from undetectable to a detectable, significant amount in the plasma of all NHPs that were injected with CBE (**Figure 4B**). We also observed a dose-dependent increase in Lyso-GL1 in liver tissue of these NHPs (**Figure 4C**). For analysis in brains, we surveyed a total of 64 brain punches to capture findings across the entirety of the NHP brain tissue. Of these 64 punches, 47 punches represent 19 grey matter regions, and the remaining 17 punches represent 7 white matter regions. Consistent with the liver data, we observed a dose-dependent increase in Lyso-GL1 in brain punches of all CBE-treated NHPs with 10 mg/kg and 30 mg/kg yielding a significant increase in Lyso-GL1 levels compared to the “No CBE” group (**Figure 4D**). Furthermore, we observed a concomitant decrease in the GCase activity of brain punches from CBE-treated NHPs (**Figure 4E**) consistent with the mechanism of action of CBE inhibiting GCase enzyme activity. We also observed this effect at the individual brain region level **(Figure S2A, B)** suggesting robust action of CBE throughout the brain tissue. As an extension of developing the CBE-induced lipid flux model in NHPs, we also performed pharmacokinetic analyses to better understand the kinetics of Lyso- GL1 (peak and clearance) in plasma with 10 mg/kg and 30 mg/kg CBE **(Figure S2C)**. We observed a similar profile of Lyso-GL1 decrease with time with both concentrations, over the course of 4 days. The increase in lipid levels in NHP brain tissues suggests target engagement with expected mechanism of action of CBE. Hence, we included CBE treatment in our pharmacology (**Figure 5**) and dose-range finding study (DRF, **Figure 6**) of SS3-GBA1 in NHPs.

**Figure 4.**
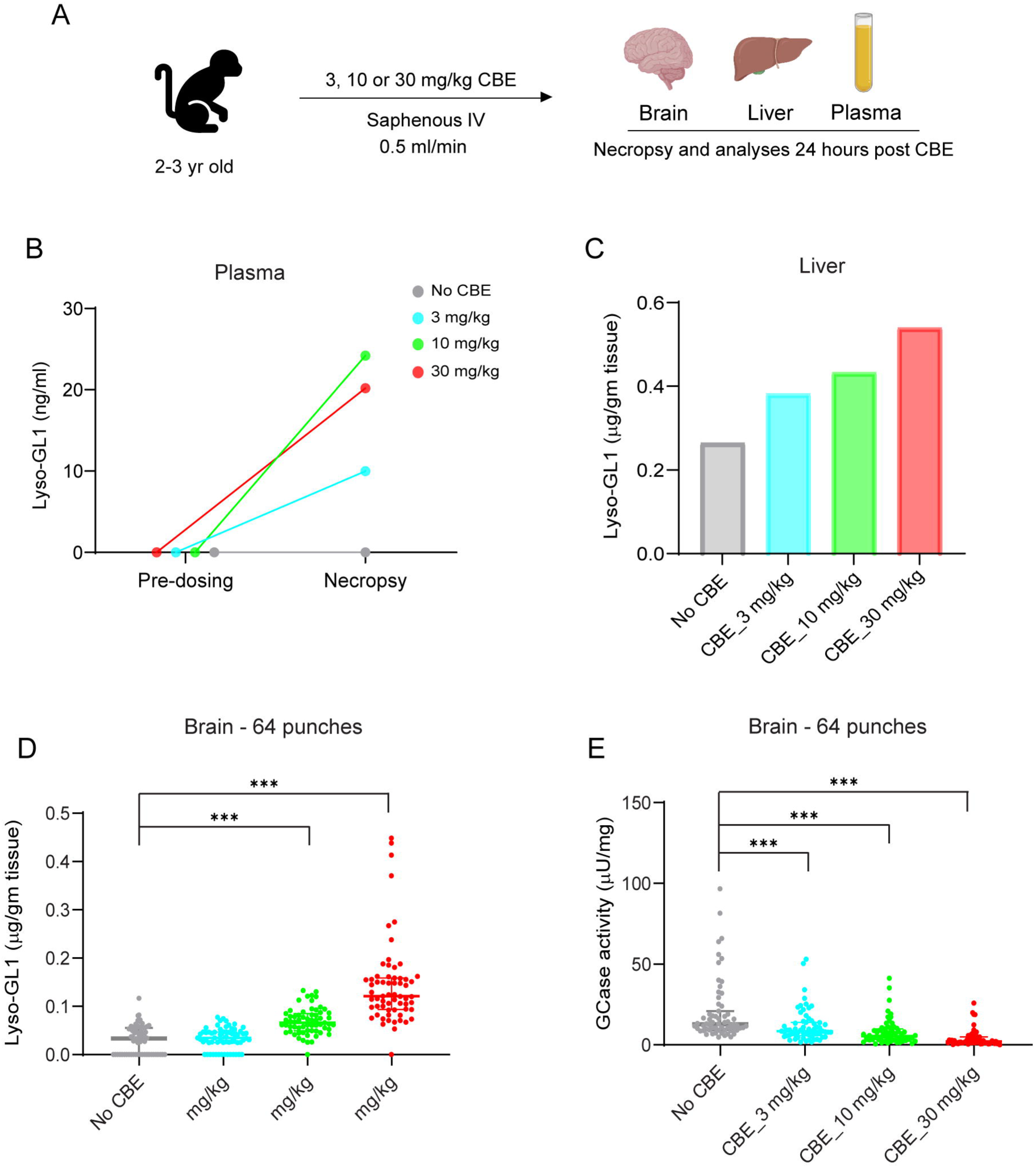
Development of CBE-induced lipid flux model in NHPs. (A) Study design: 2-3 years old cynomolgus monkeys (2-3 kg) were administered with the indicated doses of CBE (conduritol β-epoxide) by saphenous IV infusion at 0.5 ml/min, 1 animal per group. Brain, liver, and plasma samples were collected 24 hours post CBE administration for lipid analyses. (B and C) Measurement of Lyso-GL1 level in plasma samples prior to CBE dosing and 24 hours post CBE (B) and dose-dependent increase in Lyso-GL1 level in liver tissue homogenates (C) at necropsy. (D and E) Samples represent 64 brain biopsy punches encompassing 19 distinct grey matter regions and 7 distinct white matter regions. Dose-dependent increase in Lyso-GL1 (D) and concomitant decrease in GCase enzyme activity (E). 1 NHP per dose. Median with inter-quartile range across 64 punches. ***p<0.001; Two way-ANOVA with Tukey’s multiple comparisons test with all groups compared to 0 mg/kg CBE group.

**Figure 5.**
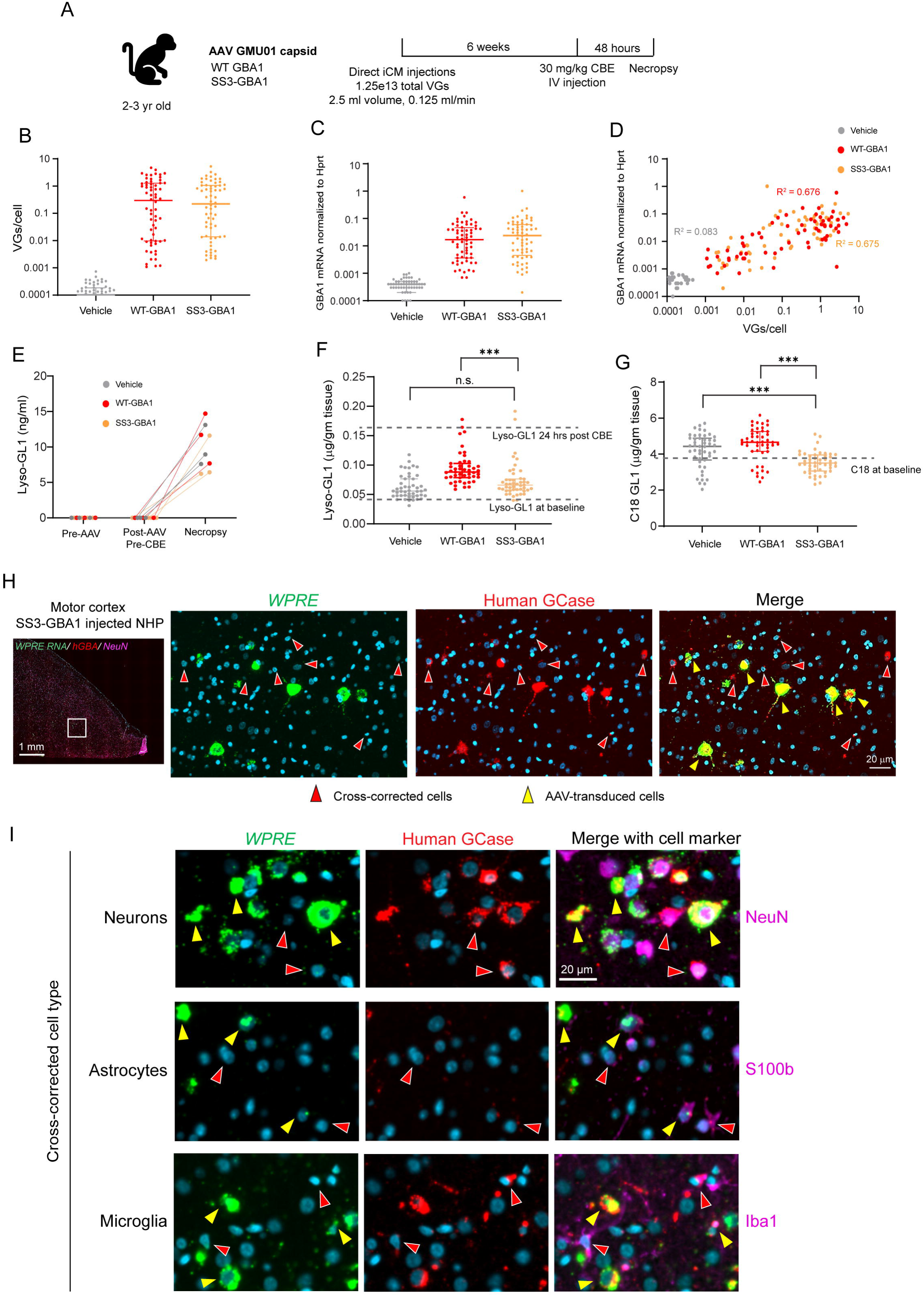
SS3-GBA1 promotes efficient lipid substrate clearance, robust secretion, and cross-correction in NHPs. (A) Study design: 2-3 years old cynomolgus monkeys (2-3kg) were dosed either with AAV GMU01-WT-GBA1 or AAV GMU01-SS3-GBA1 at 1.25e13 vector genomes per animal by direct injection to the cisterna magna (ICM) in 2.5 ml volume at 0.125 ml/min rate. 6 weeks post- dosing, 30 mg/kg of CBE was administered by IV injection 48 hours prior to necropsy. Samples represent 64 brain biopsy punches encompassing 19 distinct grey matter regions and 7 distinct white matter regions. (B and C) AAV vector genome copies determined from 64 brain biopsy punches by *Gba1* dPCR and normalized to the *Tubb3* gene copy number to obtain VG copies per cell (B). Transgene (mRNA) expression measured by GBA1 RT-dPCR normalized to endogenous *Hprt* gene (C). Median with inter-quartile range across 64 punches representing 19 grey matter and 7 white matter regions. (D) Correlation of vector genome with transgene expression (mRNA) was determined between WT-GBA1 and SS3-GBA1. Each data point is average of all NHPs for that punch. Non- parametric Spearman’s rank correlation. (E) Lyso-GL1 changes in plasma pre-AAV, post-AAV and pre-CBE, and at necropsy across all NHPs. (F and G) Lyso-GL1 level (F) and C18 GL1 level (G) across 64 brain biopsy punches. Each data point is average of all NHPs in the group for that punch. ***p<0.001; Two way-ANOVA with Tukey’s multiple comparisons test. (H and I) Multiplex fluorescent imaging assay with *in situ* hybridization for mRNA and immunohistochemistry for GBA1 protein and cell marker. Low magnification image showing stained motor cortex region of SS3-GBA1 injected NHP and zoomed-in images corresponding to the white box (H). Triplex to determine identity of cross-corrected cells using specific cell type markers, NeuN for neurons, S100b for astrocytes, and Iba1 for microglia (I). Yellow arrowheads point to AAV-transduced cells and red arrowheads point to cross-corrected cells. Scale bar is 20 microns.

**Figure 6.**
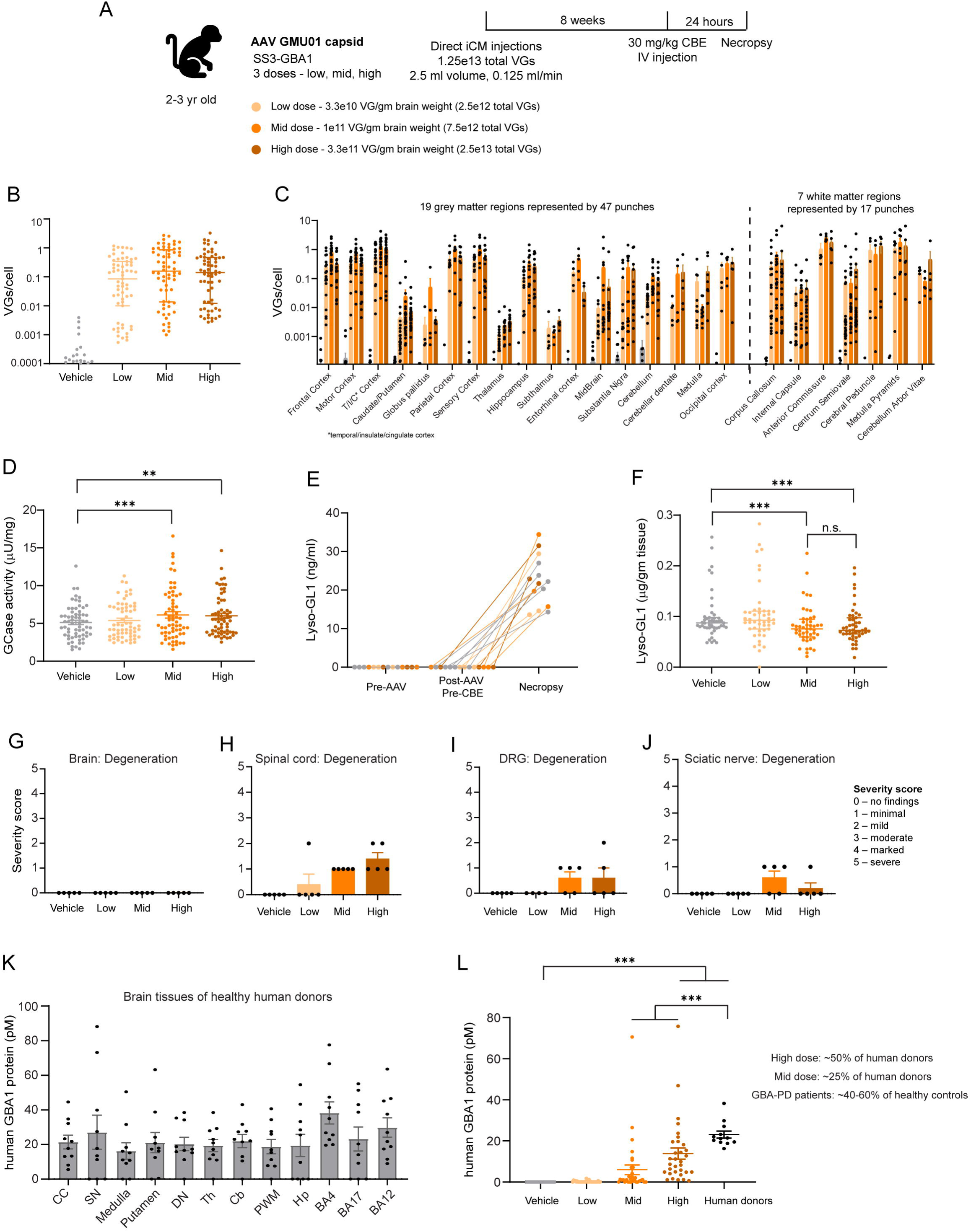
Identification of minimally efficacious dose of AAV GMU01-SS3-GBA1 in NHPs from dose-range finding study. (A) Study design: 2-3 years old cynomolgus monkeys (2-3kg) were administered with 3 different doses of AAV GMU01-SS3-GBA1 by direct injection to the cisterna magna (iCM) in 2.5 ml volume at 0.125 ml/min rate, 2.5e12 vector genomes per animal (low dose), 7.5e12 vector genomes per animal (mid dose), and 2.5e13 vector genomes per animal (high dose). 8 weeks post-dosing, 30 mg/kg of CBE was administered by IV injection 24 hours prior to necropsy. Samples represent 64 brain biopsy punches encompassing 19 distinct grey matter regions and 7 distinct white matter regions. (B and C) Assessment of vector genomes in dose-range finding study with 3 doses tested, 5 NHPs per group. Median with inter-quartile range across 64 punches representing 19 grey matter and 7 white matter regions (B). Same data shown across different brain regions (C). T/I/C: temporal/insulate/cingulate (D) GCase enzyme activity in dose-range finding study with 3 doses tested, N=5 NHPs per group. Median with inter-quartile range across 64 punches representing 19 grey matter and 7 white matter regions. **p<0.05, ***p<0.001; Two way-ANOVA with Tukey’s multiple comparisons test. (E and F) Lipidomics performed in 3 NHPs per group. Lyso-GL1 level measured in plasma (E) and from 47 brain biopsy punches from 19 grey matter regions. Median with inter-quartile range across punches. ****p<0.0001, ***p<0.001; Two way-ANOVA with Tukey’s multiple comparisons test. (G – J) Histopathological analyses in NHPs with AAV.GMU01 SS3-GBA1 dose-range finding study. Histopathological findings reported with severity scores across both central nervous system and peripheral tissues by board certified clinician. Scores reported for brain (G), spinal cord (H), DRGs (I), and sciatic nerve (J). Data are Mean ± SEM. Each dot is score for individual NHP. (K) Quantification of human GBA1 protein levels in 12 brain regions from N=10 healthy human donor brain tissues (aged 55-to-75 years old) by LC-MS. Human tissue was obtained from the NIH Neurobiobank at the University of Maryland, Baltimore, MD and the Sepulveda Research Corporation. Data are Mean ± SEM. Each data point represents brain biopsy punches from each human donor. CC: corpus callosum; SN: substantia nigra; DN: dentate nucleus; Th: Thalamus, lateral nuclear group; Cb: cerebellum; PWM: periventricular white matter; Hp: hippocampus; BA: Brodmann area. (L) Quantification of human GBA1 protein levels from 3 NHPs per group. Median with inter- quartile range across 32 punches from NHPs plotted against the 12 punches from human donors. ***p<0.0001; Two way-ANOVA with Tukey’s multiple comparisons test.

### SS3-GBA1 is safe, well tolerated and promotes efficient substrate clearance in NHPs

To test our therapeutic strategy in a large animal model, we examined biodistribution, tolerability and efficacy of SS3-GBA1 in comparison to WT GBA1 in CBE-induced lipid flux model of NHPs. The AAVs were injected via intra-cisterna magna (iCM) route, allowed to express for 6 weeks and 48 hours prior to necropsy, all NHPs were injected with 30 mg/kg CBE intravenously (**Figure 5A**).

Vector exposure (**Figure 5B**) and transgene expression of huGBA1 mRNA (**Figure 5C**) was comparable across all the 64 brain punches between WT GBA1 and SS3-GBA1. We also observed comparable correlation between transduction and mRNA levels across both AAV-treated groups (**Figure 5D**). Consistent with our previous CBE-related data in NHPs, we observed an increase in Lyso-GL1 in the plasma of all CBE-treated NHPs at necropsy (**Figure 5E**). We then performed lipidomics on all the grey matter punches to determine Lyso-GL1 and GL1 levels. Although we did not observe a significant decrease in Lyso-GL1 levels in NHPs treated with either AAV when compared to vehicle injected NHPs, Lyso-GL1 was still significantly lower in SS3-GBA1 treated NHPs compared to WT GBA1 treated NHPs (**Figure 5F**). Further, examination of individual GL1 species revealed that SS3-GBA1 significantly reduced accumulation of C18 GL1, a predominant isotype of GL1 in the brain (*21, 33*), not only in comparison to the vehicle group but also the WT GBA1 treated NHPs (**Figure 5G**). Importantly, AAV.GMU01 SS3-GBA1 treatment lowers the accumulated C18 GL1 to physiological levels (grey line indicating physiological C18 GL1 levels from our pilot CBE study in NHPs).

One explanation for why we did not observe a significant reduction in Lyso-GL1 with SS3-GBA1 is that the 48-hour window between CBE-dosing and necropsy was likely too long to capture the peak Lyso-GL1 accumulation. Our pilot CBE study in NHPs was analyzed 24 hours post CBE dosing and by 48 hours Lyso-GL1 in vehicle or untreated NHPs drops down 3-fold (0.15 at 24 hours and 0.05 at 48 hours, median of uninjected **Figure 4D** and vehicle **Figure 5E**). Consistent with this speculation, we indeed observed robust clearance of Lyso-GL1 with SS3-GBA1 in our DRF study (**Figure 6**) wherein we performed necropsy 24 hours post CBE dosing. Nevertheless, since Lyso-GL1 is a result of deacetylation of GL-1 by acid ceramidase (*34*), reduction in GL-1 lipids is indicative of reduced levels of Lyso-GL1 as well. Together, these findings suggest that SS3-GBA1 is not only efficacious in rodents but also in large animal models.

We also performed histopathological analyses in the brain, spinal cord, sciatic nerve and dorsal root ganglion (DRGs), to examine any tissue degeneration associated with AAV-dosing. While both WT GBA1 and SS3-GBA1 were well-tolerated with minimal to mild findings, SS3-GBA1 was better tolerated than WT GBA1 in some tissues such as spinal cord and sciatic nerve (**Figure S3A-D**). Given the potent secretory nature of SS3-GBA1, it is likely that SS3-GBA1 has more widespread distribution with overall less abundant protein levels in each tissue/region which may explain the improved tolerability for SS3-GBA1 injected NHPs, compared to WT GBA1.

### Robust secretion and cross-correction of SS3-GBA1 in NHP brain tissues

To generate clear evidence of SS3-GBA1 cross-correction, we developed a fluorescent multiplex ISH/IHC (*in situ* hybridization for mRNA and immunohistochemistry for protein distribution) assay to capture both AAV transduced cells and protein biodistribution with multiplex imaging in the same tissue section. We performed *in situ* hybridization to identify cells transduced by the AAV vector, expressing the transgene and immunohistochemistry with an anti-human GBA1 antibody to stain for human GCase protein. Consistent with our results in rodent studies, we observed clear examples of cross-corrected cells throughout the NHP brain tissue (**Figure 5H**). Representative high magnification images from the motor cortex show AAV-transduced cells (yellow arrows in merge that are positive for mRNA in green and human GCase protein in red) and several non-transduced cells positive for human GCase protein (red arrows).

We next determined the identity of cross-corrected cells, by adding in immunostaining for cell markers in our multiplex assay. We observed not only neurons but also astrocytes and microglia able to uptake the secreted huGBA1 protein, indicating robust cross-correction of multiple cell types within the brain tissue (**Figure 5I**). This is an important finding given the ubiquitous nature of *GBA1* across different cell types in the brain. SS3-GBA1 cross-correcting within multiple cell types is likely a key factor in driving therapeutic efficacy, which explains the robust lipid clearance we observe across species.

Taken together, our data reveal that AAV.GMU01 SS3-GBA1 is well tolerated and superior to WT GBA1 in terms of efficacy not only in mice but also in NHPs. SS3-GBA1 is robustly secreted, cross-corrected and promotes lipid clearance across species. Furthermore, SS3- GBA1 cross-corrects multiple cell types within the brain which lays the foundation for a large therapeutic footprint, driving efficacy of our gene therapy strategy.

### Dose-range finding study (DRF) in NHPs identifies minimally efficacious dose of AAV.GMU01 SS3-GBA1

Based on all the data thus far, we nominated SS3-GBA1 as our lead candidate and performed a dose-range finding study in NHPs to evaluate biodistribution, tolerability and efficacy of SS3- GBA1 at three different doses of the AAV vector – low (3.3e10 VG/gm brain weight), mid (1e11 VG/gm brain weight) and high (3.3 e11 VG/gm brain weight) (**Figure 6A**). Given our lipidomics data in our NHP pharmacology study wherein we failed to observe clearance of Lyso-GL1 in SS3-GBA1 injected NHPs when compared to vehicle group, we decided to reduce the time window between CBE dosing and necropsy from 48 hours to 24 hours, to evaluate efficacy of SS3-GBA1 in the context of peak accumulation of Lyso-GL1.

Across all three doses, we observed a broad brain distribution of the AAVs both in grey matter as well as white matter brain regions (**Figure 6B, C**). We also observed significant increase in GCase activity across the 64 brain punches, with the mid and high dose yielding significantly higher GCase activity than vehicle injected group (**Figure 6D**). Before performing lipidomics in brain grey matter tissues, we first determined Lyso-GL1 levels in the plasma of all NHPs at three timepoints: pre-AAV, post AAV pre-CBE and post-CBE necropsy timepoints. Lyso-GL1 levels in plasma increased after CBE administration, as measured in previous study (**Figure 6E**). Lipidomic analyses in brain punches revealed a significant reduction in Lyso-GL1 in mid and high-dose groups, indicating effective clearance of accumulated Lyso-GL1 by SS3-GBA1 (**Figure 6F**). Importantly, both mid and high-dose had similar efficacy levels indicating a maximal effect on Lyso-GL1 and further establishing our mid dose of 1e11 VG/gm brain weight as the minimally efficacious dose in the context of lipid clearance. We also performed histopathological analyses across all three doses and consistent with our pharmacology study, we only observed minimal to mild microscopic findings across all tissues surveyed (**Figure 6G-J**, **Figure S2E, F)**. In summary, we conclude that a single administration of SS3-GBA1 via iCM injection has a broad CNS distribution, is safe, well tolerated, and is efficacious.

### AAV.GMU01 SS3-GBA1 dosing results in huGBA1 protein levels that is therapeutically beneficial for GBA-PD patients

GBA-PD patients are carriers of heterozygous mutations in their *GBA1* gene, resulting in haploinsufficiency with reduced enzymatic activity, reduced GCase protein levels, elevated lipids and increased α-synuclein (*14, 15, 20, 21, 35, 36*). Given the strong correlation between reduced GCase protein levels with disease severity and disease progression, we next wanted to determine if our AAV gene therapy strategy can replenish the deficit in human GCase protein levels that has previously been reported in GBA-PD brain tissue samples. To this end, we first procured post-mortem brain tissues of healthy human donors between the ages of 55 to 75 years-old, which is the most common age of onset for PD. We developed a proteomics mass spectrometry assay to specifically detect human GCase protein in these human tissues as well as in NHPs injected with our AAV vectors. In human brain tissues, we selected 12 different brain regions that largely represented the different functional regions and measured human GCase protein levels in these regions across 10 healthy human donor brains (**Figure 6K**). In NHPs, we surveyed 32 different brain punches, and we observed a dose-dependent increase in human GCase protein levels in NHPs, but overall lower protein levels compared to human brain samples (**Figure 6L**). In the high dose NHP group, our AAV dosing yielded human GCase protein levels that were about ∼50% of the human brain values. In the mid-dose NHP group, AAV dosing yielded human GCase protein levels that were about ∼25% of the human brain values. Given that GBA-PD patients typically carry one functional copy of the *GBA1* gene, replenishing 50% of control human GCase protein levels in these patients would be sufficient to restore GCase level to healthy physiological levels. Multiple studies examining GCase protein levels and enzymatic activity in GBA-PD patients have now reported that these patients have a deficit of ∼40-60% of GCase protein levels when compared to healthy controls (*14, 37*). Our high dose resulted in ∼50% of healthy control human GCase protein amount, indicating potentially clinically relevant restoration of GCase protein levels in GBA-PD patients. Hence, we conclude that a one-time iCM dosing of AAV.GMU01 SS3-GBA1 results in potentially therapeutically meaningful human GCase protein levels that would replenish the deficit of GCase protein seen in GBA-PD patients, correcting the underlying biology in these patients.

### Single dose of AAV.GMU01 SS3-GBA1 results in long-term efficacy and durability in *Gba^D409V/+^ mice*

All studies thus far were evaluated for efficacy either 4 or 6-weeks post AAV dosing. To investigate durability and efficacy of our gene therapy strategy on longer time scales, we performed a long-term study in the *Gba^D409V/+^*mice (*30–32*) wherein we injected AAV.GMU01 SS3-GBA1 via bilateral ICV and cohorts of mice were taken down at either 3 months, 6 months or 9 months post-AAV injection (**Figure 7A**). 24 hours prior to necropsy, all cohorts also received 100 mg/kg CBE to induce lipid increase, which allowed us to assess efficacy via lipid clearance. To demonstrate cross-correction of our secreted payload, we microdissected the brain into 4 regions – cortex, sub-cortical (proximal to site of injection), and hindbrain, cerebellum (distal to site of injection) (**Figure 7B**) and performed lipidomics to survey lipid clearance over longer duration. Consistent with bilateral ICV injections, we observed higher VGs in the cortex and sub-cortical regions and an order of magnitude lower VGs in the cerebellum and hindbrain (**Figure 7C-F**). Despite large differences in vector exposure in proximal and distal sites, we observed significant decrease in Lyso-GL1 levels across all regions, across all time-points (**Figure 7C-F**) indicating secreted GCase distributing throughout the brain. We also observed the robust clearance of lipids in plasma of SS3-GBA1 injected mice, across all time points (**Figure 7G**). These data indicate that a one-time administration of SS3-GBA1 is sufficient to actively promote lipid clearance over longer time periods, demonstrating sustained efficacy over time.

**Figure 7.**
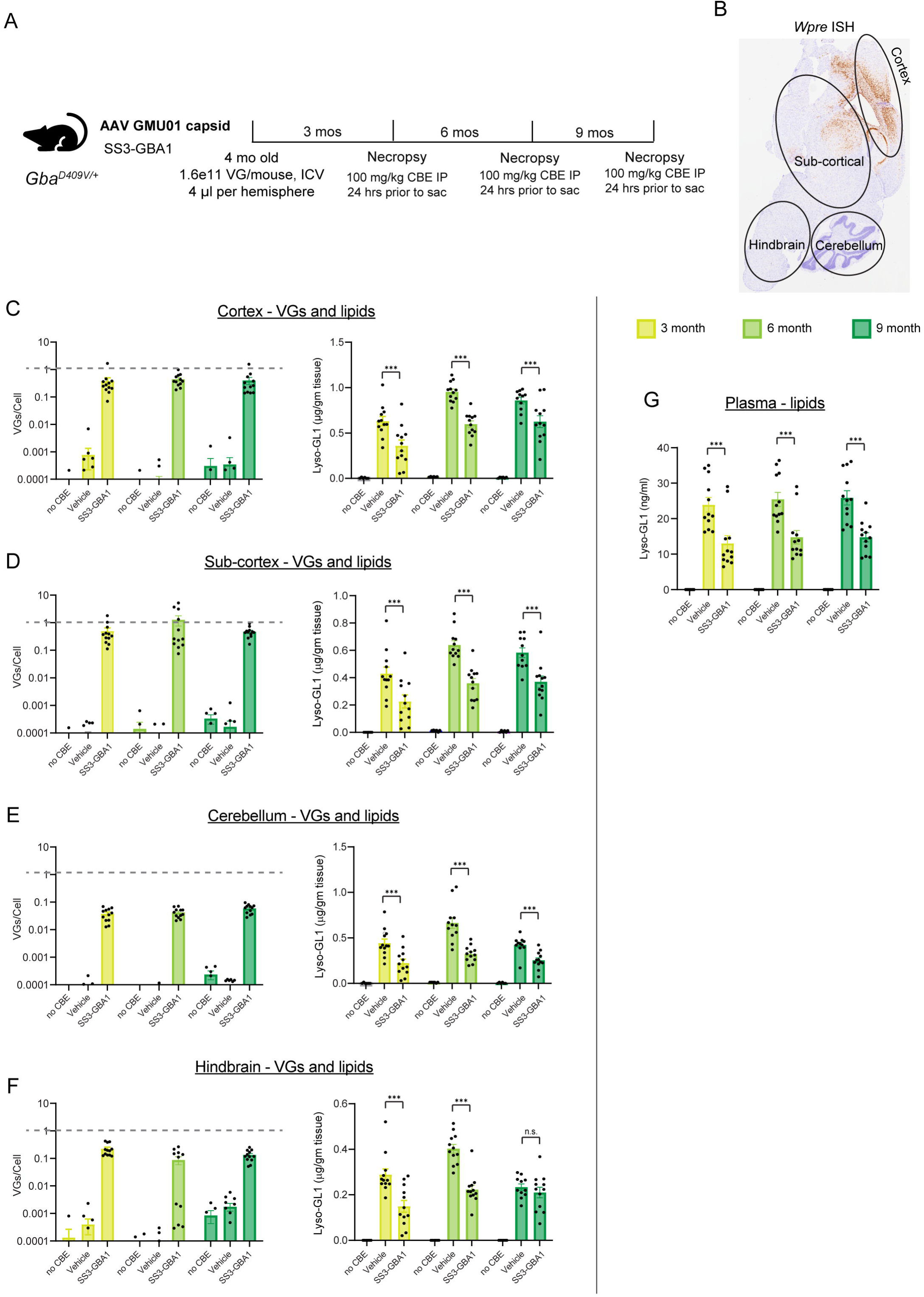
Long-term efficacy and durability with AAV GMU01-SS3-GBA1 in *Gba^D409V/+^* mice. (A) Study design: AAV GMU01-SS3-GBA1 was administered by bilateral ICV to 4-month-old *Gba^D409V/+^*mice at 1.6e11 vector genomes per animal and 4μl per hemisphere. 6-12 animals per group. Brain, plasma, and CSF were analyzed 3-month, 6-month, and 9-month post-injection. CBE (conduritol β-epoxide) was administered at 100 mg/kg 24 hours prior to each timed necropsy. (B) Representative image of *in situ* hybridization to WPRE mRNA from the analyzed sagittal sections. (C – F) Vector genome assessment of the longitudinal pharmacology study with in-life duration of 3-month, 6-month, and 9-month post AAV-dosing. AAV vector genome copies (left) and Lyso- GL1 lipid clearance (right) in cortex (C), sub-cortex (D), cerebellum (E), and hindbrain (F) across all mice in the study. 6 animals for No CBE group, 12 animals for Vehicle group, and 12 animals for SS3-GBA1 group. Data are Mean ± SEM. ***p<0.0001; Two way-ANOVA with Tukey’s multiple comparisons test. (G) Lyso-GL1 lipid clearance in plasma of SS3-GBA1 injected mice. Data is Mean ± SEM. ***P<0.0001; Two way-ANOVA with Tukey’s multiple comparisons test.

### IV administration of AAV.GMU01 SS3-GBA1 effectively targets peripheral tissues affected in Gaucher Disease

Given that bi-allelic mutations in *GBA1* are causative for Gaucher Disease (GD), a rare lysosomal storage disease, we hypothesized that our lead candidate could serve as a disease modifying therapy for GD Type 1, when administered intravenously to target all the peripheral tissues implicated in GD Type 1. To test this, we investigated the biodistribution and efficacy of AAV.GMU01 SS3-GBA1 when administered via IV dosing. We injected either vehicle or AAV.GMU01-SS3-GBA1 in 3-month-old mice (4e13 VG per kg), allowed AAV expression for 4 weeks followed by 100 mg/kg injection of CBE, 24 hours prior to necropsy (**Figure 8A**).

**Figure 8.**
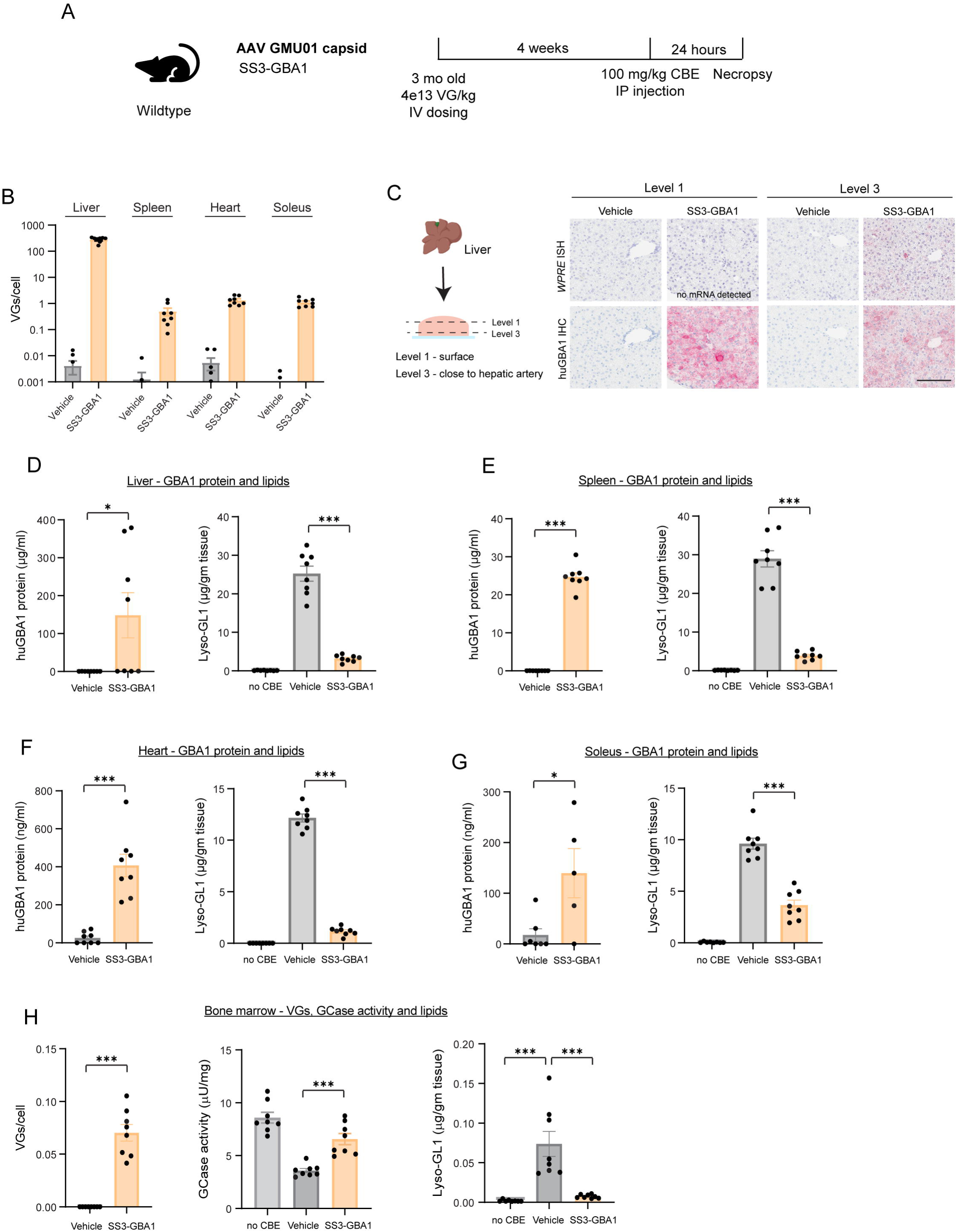
Effective targeting of peripheral tissues affected in Gaucher disease by IV administration of AAV GMU01-SS3-GBA1. (A) Study design: 3-month-old C57/BL6 mice were injected with 4e13 VG/kg of vehicle or AAV GMU01-SS3-GBA1 intravenously. AAVs were expressed for 4 weeks followed by 100mg/kg of CBE IP injection 24 hours prior to necropsy. (B) Vector genome assessed in different visceral organs such as liver, spleen, heart, and soleus muscle. 8 animals per group. Data is Mean ± SEM. (C) Schematic view of liver sectioning strategy and representative images demonstrating robust *GBA1* secretion. Vector biodistribution is shown with *in situ* hybridization to WPRE (top panels) and GBA1 protein expression is shown with huGBA1 immunohistochemistry after 4 weeks of expression. (D – G) Quantification of human GBA1 protein level by ELISA and Lyso-GL1 lipid clearance by LC-MS in liver (D), spleen (E), heart (F), and soleus muscle (G). 8 animals per group, No CBE group in the graphs shown is control mice that did not receive any AAV vector or CBE injection. Data are Mean ± SEM. *p<0.05, ***p<0.0001; unpaired Student’s t-test (human GBA1 protein) and ***p<0.0001; One way-ANOVA with Dunnett’s multiple comparisons test (Lyso-GL1). All groups compared to vehicle group. (H) Evaluation of vector exposure, GCase enzyme activity, and Lyso-GL1 lipid clearance from bone marrow. 8 animals per group and No CBE group in the graphs shown is control mice that did not receive any vector or CBE injection. Data are Mean ± SEM. ***p<0.0001; unpaired Student’s t-test (VGs/cell) and ***p<0.001; One way-ANOVA with Tukey’s multiple comparisons test (GCase activity and Lyso-GL1).

We first examined liver, spleen, heart and soleus muscle for vector exposure and lipid clearance. Consistent with IV dosing, we observed high vector distribution within the liver (**Figure 8B**). We also observed robust cross-correction within the liver as seen by *in* situ hybridization to detect transgene mRNA and immunohistochemistry to detect human GCase protein (**Figure 8C**). We observed higher mRNA transgene proximal to the hepatic artery resulting in widespread protein biodistribution. Further away from the artery, we obtained little to no AAV vector transduction, yet we observed broad protein biodistribution indicating cross-correction of our SS3-GBA1 payload. Consistent with this, we observed robust lipid clearance across all tissues we surveyed (liver, spleen and muscles such as heart and soleus) (**Figure 8D-G**). To specifically measure huGBA1 protein levels in bulk tissues, we developed an in-house sandwich ELISA **(Figure S3A-C)**. Although we observed varying amounts of huGBA1 protein across these different tissues, our lipidomics data suggest that even low levels of functional GCase can be highly effective in driving lipid clearance. For example, although huGBA1 protein levels within the spleen was an order of magnitude lower than that in heart and soleus muscle tissues, all tissues had comparable clearance of accumulated lipids. These findings prompted us to perform lipidomics on other major peripheral tissues as well as other muscles. Indeed, across all tissues we surveyed – lung, kidney, diaphragm, quadriceps and gastrocnemius, we observed significant lipid reduction **(Figure S3D-H)**, suggesting broad biodistribution and effective cross-correction of AAV.GMU01 SS3-GBA1.

The current standard of care for GD patients typically involves enzyme replacement therapy (ERT) which involves bi-weekly infusions of the recombinant GBA1 protein. While this treatment manages symptoms for most people, it does not prevent disease progression in tissues that are not targeted by the ERT. GD patients typically present with “Gaucher cells” infiltrating the bone marrow and only high doses of therapy seem to target these cells, albeit with limited efficacy in some patients. Bone marrow (*38–40*). To determine if our gene therapy strategy targets cells in the bone marrow, we isolated these cells from AAV-injected mice and evaluated vector exposure, GCase activity and lipid clearance. Despite low VGs, we observed a significant increase in GCase enzyme activity and significant decrease in lipids accumulated compared to vehicle injected mice (**Figure 8H**). These data indicate effective huGBA1 protein distribution within the bone marrow contributing to its functional efficacy in promoting lipid clearance. Collectively, our IV data position our therapeutic strategy superior to the conventional ERT given its broad therapeutic footprint and its ability to target tissues that are typically impenetrable by ERT.

## Discussion

*GBA1* or GCase is a unique lysosomal enzyme that is involved in two distinct diseases – Gaucher disease which is the most prevalent lysosomal storage disease and GBA-related Parkinson’s disease affecting 0.5-1 million people worldwide (*8*). Despite the different clinical trajectories of these two diseases, recent evidence in PD research (*41, 42*) indicates that these diseases converge on lysosome dysfunction as a potential mechanistic basis of the underlying disease biology. Here, we propose a gene therapy strategy to treat both GBA-PD and GD with a single-administration of our gene therapy product. Using a combination of studies across species, we demonstrate safety, tolerability and efficacy of our strategy.

Given the different tissues affected in GD and GBA-PD, a key aspect to ensure efficacy of a therapeutic candidate across the two diseases is targeting multiple tissues effectively, generating a broad biodistribution profile. “Cross-correction” is a phenomenon of secreted enzymes taken up by neighboring cells that are deficient for the functional enzyme. Cross-correction, is thus, one way to enable broad biodistribution of proteins both within and across tissues. We applied this principle to our payload and engineered it to be readily secretable. We present multiple lines of evidence that demonstrate robust cross-correction of engineered *GBA1* transgene: (i) fluorescent imaging to examine AAV transduced cells and non-transduced cells alongside human GCase protein, (ii) human GCase protein quantification data in tissues with little to no AAV vector transduction, (iii) lipidomics showing substrate clearance on microdissected brain regions distal to injection site. Thus, the combination of our novel capsid and engineered payload reveals effective tissue targeting and broad biodistribution of huGBA1 protein.

To identify the lead candidate across our GCase variants, we developed a CBE-induced lipid-flux model in both mice and NHPs. We identified a working CBE regime that allows us to query the efficacy of our lead candidate via lipidomics performed on rodent and NHP tissues. The CBE-dosing regimen identified in mice and NHPs worked reproducibly and consistently across our studies, and is an invaluable tool for investigating efficacy of therapeutic strategies aimed at GBA1-related diseases. Efficacy data from both mice and NHPs demonstrate SS3-GBA1, our lead candidate to be superior to WT-GBA1. Additionally, SS3-GBA1 in NHPs resulted in lower histopathological scores than WT-GBA1.

Using the CBE-model in NHPs, we have also identified the minimally efficacious dose with a single administration of AAV.GMU01 SS3-GBA1. We obtained similar reduction in Lyso-GL1 lipid levels with both our mid- and high-dose of AAV (1e11 VG/gm brain weight and 3.3e11VG/gm brain weight). Despite the high dose resulting in significantly higher huGBA1 protein amount in NHP brain tissues than the mid-dose, we failed to observe an even higher clearance of Lyso-GL1 in the high-dose compared to the mid-dose. One likely reason for this could be the rate-limiting amount of co-factors that GCase protein requires for its role in hydrolyzing the lipids. Saposin C is indeed a co-factor that has been shown to interact with GCase and is necessary for its action on lipid breakdown (*43, 44*). Nevertheless, our data suggests that we have a clinically feasible dose resulting in therapeutic efficacy.

We also performed proteomic analyses on NHPs injected with SS3-GBA1 and compared the human GCase protein levels in these NHP brain tissues with that found in healthy human donor brains aged 55-70 years old. By estimating the physiological protein levels in human brain tissues, we can establish a reference point to determine whether our gene therapy approach produces human GCase protein levels that might be clinically relevant. Our data reveal that we can replenish about ∼25% and ∼50% human GCase protein levels in our mid- and high-dose groups respectively. Given that GBA-PD patients have a deficit of ∼40-60% of GCase protein amount (*14, 37*), our data suggest that the combination of our capsid and payload is indeed able to produce therapeutically beneficial amount of GCase protein levels that are required for GBA-PD patients. Furthermore, our longitudinal pharmacology study in *Gba^D409V/+^* mice indicate that single administration of AAV.GMU01 SS3-GBA1 results in long-term efficacy and durability.

While GBA-PD is a disease of the central nervous system (CNS), the most common form of Gaucher Disease (Type 1) is largely peripheral. To demonstrate that our gene therapy product could also be beneficial for GD Type 1 patients, we performed efficacy studies in mice with IV dosing. Our cross-correction strategy resulted in broad tissue targeting with robust lipid clearance across major peripheral organs such as liver, spleen and muscles. Although there are multiple therapies currently available for GD patients, the major clinical unmet need for these patients are two-fold: (i) current treatments have little to no penetration into the bone and bone marrow cells and hence most patients present with severe bone complications over time that are largely unaddressed with most treatments (ii) GD Type 3 or neuronopathic GD patients who not only have peripheral tissue symptoms but also severe neurological symptoms and none of the current treatments address the rapidly progressing CNS component of the disease in these patients (*2*). A remarkable aspect of our strategy is that we present preclinical data that addresses both these clinical needs - effective targeting of and lipid clearance in bone marrow as well as engineered payload that drives efficacy in multiple brain regions with intra-CSF dosing, which targets the CNS component of the disease. Although this means the route of administration for Type 1 GD patients would be IV and for Type 3 GD patients it would be intra-CSF, these seem appropriate given the differential symptoms between these patient subtypes.

Finally, a compelling feature of our gene therapy strategy is the cross-correction we observe not only in neurons but also in other cell types in the brain such as astrocytes and microglia. This finding alone has major implications for the underlying disease biology in not only GBA-PD but also idiopathic PD. Although neuroinflammation in idiopathic PD patients is well established (*45–48*), only recent studies have uncovered the role of *GBA1* in astrocytes and microglia in regulating tissue homeostasis and neuroinflammation via astrogliosis and microglial activation. For example, astrocytes lacking GCase or harboring loss-of-function mutations in *GBA1* exhibit astrogliosis with impaired lysosomal and proteasomal activity that are in turn implicated in mitophagy and clearance of aggregated α-synuclein (*33, 49, 50*). In the same vein, lack of GCase in microglia makes the neurons more susceptible to toxic insults and neurodegeneration (*51*); and *GBA1* rescue in microglia alone prolonged survival in neuropathic GD mice (*52*) implicating a key role for microglial *GBA1* in neuronal health and survival. Interestingly, a recent imaging study in *GBA1* mutation carriers without PD symptoms or diagnosis also revealed an increase in microglial activation (*53*), suggesting that the effects of loss of GCase in microglia and astrocytes likely precedes downstream effects seen in neurons with lipid accumulation and α-synuclein accumulation. Thus, our ability to cross-correct multiple cell types is an important aspect of our strategy in driving therapeutic and clinical benefit.

Collectively, our results lay the foundation for a clinically relevant and well-tolerated gene therapy approach with our novel capsid and engineered payload. A single-administration of AAV.GMU01 SS3-GBA1 robustly cross-corrects multiple cell types, is efficacious, well-tolerated with no adverse effects, offering a compelling therapeutic strategy that could be highly beneficial for the GBA-PD and GD patients.

## Materials and Methods

### Animal care and use

All experiments were conducted in AAALAC accredited institutions. Animals were handled in accordance with the rules and regulations of the IACUC, in compliance with the Animal Welfare Act, and adhered to principles stated in the Guide for the Care and Use of Laboratory Animals. All effort to minimize pain and distress were ensured in these purpose-bred animals.

### Rodent

*Gba^D409V^* mouse model (The Jackson Laboratory, stock #415294, B6. *Gba^D409V/+^*) was used for CBE-dosing and long-term efficacy studies. 3-4 month old wildtype, heterozygous, and homozygous littermates were used for analyses.

### Non-human primates (NHP)

Only purpose-bred naïve cynomolgus NHPs (2-3 year old and 2- 3kg) were used in studies. NHPs were prescreened for AAV neutralizing antibodies, and seronegative animals were selected for the studies.

### Vector/plasmid design

#### CBA-WT GBA-WPRE-bGH

Functional wild type human *GBA1* gene (NM_000148.2) was flanked by AAV2 inverted terminal repeats (ITRs) under the control of the human CBA promoter (CMV enhancer, chicken β-actin promoter, and a chicken β-actin/rabbit β-globin hybrid intron), included an engineered Woodchuck Hepatitis Virus Posttranscriptional Regulatory Element (WPRE) to enhance GBA expression in transduced cells and the bovine growth hormone (bGH), polyA, and termination signal.

#### Engineered *GBA1* variants

The N-terminal signal sequence of human GBA1 sequence was replaced by signal sequences from other secretory proteins. Rest of the sequence in the constructs were identical to WT-GBA1 (NP_000148.2).

|  | Signal peptide sequence |
| --- | --- |
| WT-GBA1 | MEFSSPSREECPKPLSRVSIMAGSLTGLLLLQAVSWASG |
| SS1-GBA1 | MASLWLLSCFSLVGAAFG |
| SS2-GBA1 | MFQTGGLIVFYGLLAQTMA |
| SS3-GBA1 | MLLLLLLLGLRLQLSLG |
| SS4-GBA1 | MTILFLTMVISYFGCMKA |

#### In silico analyses

To generate *GBA1* variants that are readily secretable, we used two bioinformatic tools: SignalP 6.0 (*54*) and DeepLoc 1.0 (*55*). SignalP 6.0 is a machine language model that predicts/detects signal peptides in protein sequences and estimates probability of signal peptide getting cleaved. DeepLoc 1.0 is a deep neural network that predicts localization of proteins within cells with a probability score for each localization such as lysosomal, mitochondrial, extracellular.

The N-terminal signal peptide of human GBA1 protein was replaced with signal peptide of secreted proteins and this new protein sequence was then inputted into both the tools to query for signal peptide cleavage probability and predict localization in cells. Of all secreted proteins examined, 4 top candidates with highest probability of extracellular localization were chosen which we termed as signal sequence (SS), SS1 through SS4. Probabilities of signal peptide cleavage and subcellular localization of these 4 candidates are presented in Table S1.

### Culturing and transfection of HEK293T cell line

HEK293T (ATCC, CRL-3216) cells were maintained in DMEM (Gibco, 11965092) media supplemented with 10% (vol/vol) FBS (Gibco, A3160601) under standard cell culture conditions, 37°C in 5% (vol/vol) CO_2_. Cells were transfected with Lipofetamine3000 (Invitrogen, L3000015) by following the manufacturer’s protocol. Plasmids were allowed to express for 48 hours before downstream analyses.

### Immunocytochemistry (ICC) and imaging

Immunocytochemistry was performed on HEK293T cells where 15,000 cells per well were seeded in 96-well microplate (Corning, 353219) and transfected with plasmid described above. Briefly, Cells were incubated with 50nM of MDW933 probe (Sanofi compound management) in cell culture media for 2 hours at 37°C to label active GCase and followed by 4% (vol/vol) paraformaldehyde (Thermofisher, J19943-K2) fixation for 15 minutes at room temperature. Permeabilization and non-specific binding blocking were achieved by 2 hours incubation in PBS solution containing 10% (vol/vol) normal donkey serum (JacksonImmuno Research, 017000121), 1% (vol/vol) BSA (Sigma, A7030), and 0.3% (vol/vol) Trition X-100 (Sigma, T9284) at room temperature. Cells were stained with anti-Lamp1 (1:200, CST, 9091) primary antibody in PBS supplemented with 5% (vol/vol) normal donkey serum, 1% (vol/vol) BSA, and 0.3% (vol/vol) Triton X-100 overnight at 4°C. Then, Cells were washed in PBS and incubated in anti-rabbit Alexa Fluor Plus 647 (1:1000, Invitrogen, A32795) secondary antibody prepared in PBS supplemented with 5% (vol/vol) normal donkey serum, 1% (vol/vol) BSA, and 0.3% (vol/vol) Triton X-100 for 1 hour at room temperature. Nuclei was counter stained with Hoechst 34580 (1µg/mL, Invitrogen, H21486). Immunofluorescence images were acquired on a PerkinElmer Opera Phenix, and spot analysis was performed with Harmony software.

### Administration of Conduritol ß Epoxide (CBE)

CBE (Sigma-Aldrich, 234599) was reconstituted with sterile saline to a concentration of 10mg/mL just prior to use. For rodents, mice were injected with 100mg/kg of CBE at 10mL/kg by intraperitoneal (IP) injection using 27 1/2-gauge needle 24 hours prior to necropsy. For non-human primates (NHPs), single dose of CBE was given to 2-4 years old cynomolgous monkeys by intravenous (bolus) infusion to the saphenous vein at 0.5mL/minutes rate 24-48 hours prior to necropsy.

### AAV manufacturing

AAV vectors were produced using the transient triple transfection method as previously described (*56*). Briefly, HEK293 cells were transfected using three plasmids (AAV ITR vector plasmid, AAV rep/cap helper plasmid, and pAd helper plasmid, (pAd Helper, Stratagene/Agilent Technologies, Santa Clara, CA). AAV vectors were purified by affinity column chromatography (POROS AAV9 Capture Select, Life Technologies) as previously described (*56*) followed by cesium chloride gradient sedimentation purification to enrich for full AAV vector genome containing capsids (*57*). For some vector preparations, purification was achieved using a combination of two rounds of cesium chloride gradient sedimentation purification. Vectors were tittered using a ddPCR assay with primer and probes targeted to the BGH sequence in the AAV vector genome. Vectors were formulated in phosphate buffer + 0.0015% F-68.

### Bilateral intracerebroventricular (ICV) injection

4-month-old C57BL/6 or mice were anesthetized with isoflurane and received bilateral injections into the lateral ventricles (A–P: −0.4; M–L: ±1.0; D–V: −2.7) with 4-5μL per site of formulation buffer or AAV.GMU01 WT GBA1 and variants using a 10-μL Hamilton syringe at a rate of 1 μL/min. Following the procedure, the injected mice were monitored, and supportive care was given for recovery.

### Intra-cisterna magna (ICM) injection

Cynomolgous monkeys were fasted overnight (at least 8 hours) prior to dosing procedure. The animals were anesthetized and positioned in lateral Trendelenburg position with an angle of approximately 15-20 degrees. The dosing setup was primed with formulation buffer prior to dosing and fluoroscopy imaging was used to guide the surgeon during the dosing. As a single step, a volume of 2.5mL of either formulation buffer or AAV.GMU01 GBA1 vector at 1.25e13 VGs per vector was administered at a rate of 0.125mL/minute. A flush volume of 0.250mL was given at the end of the dosing and the needle was left in place for at least 1-3 minutes before removal. Following the procedure, veterinary care was given for recovery.

### Intravenous (IV) injection

3-month-old C57B BL/6J mice were injected with 100 µl of formulation buffer or AAV.GMU01 WT GBA1 or AAV.GMU01 SS3 GBA1 vectors to the lateral tail vein using a 30-gauge needle.

### Tissue homogenization

NHP and rodent tissues were homogenized in ice cold TE buffer, pH 7.4 (Fisher scientific, BP2476500) either in 1.4mm ceramic bead tubes (Fisher scientific, 15-340-153) or 2.8mm ceramic bead tubes (Fisher scientific, 15-340-154) with 6.5mm ceramic beads (OMNI International, 19-682) using Omni Bead Ruptor 12 (OMNI International). Following homogenization, tissue homogenate aliquots were made for all the subsequent assays.

### Protein extraction

TE buffer, pH 7.4 (Fisher scientific, BP2476500) supplemented with Nonidet P-40 (Thermo scientific, 28324) and halt protease inhibitor cocktail (Thermo scientific, 87786) was added to collected tissue homogenates to 0.1% NP-40 final concentration. The homogenates were allowed to mix/solubilize at 4°C for 30 minutes on a tube revolver rotator (Thermo scientific, 88881001) and followed by centrifugation at 4°C and 18,000 *x g* for 10 minutes. Clear supernatants were collected and transferred into 1.5mL eppendorf tubes (Eppendorf, 22363204).

### Protein quantification

Total protein concentration was determined by BCA (bicinchoninic acid) protein assay kit (Thermo Scientific, 23225). Colorimetric detection was done by measuring absorbance at 562 nm with Molecular Devices SpectraMax 340PC 384 96-well microtiter plate reader and SoftMax Pro version 5.4.4 software.

### Glucosylceramidase activity assay

#### In-house GCase activity assay

Recombinant human GBA (Cerezyme, Sanofi) standard was prepared using lysis buffer composed of 50mM K2HPO4 dibasic (Sigma, P8584), 50mM K2HPO4 monobasic (Sigma, P8709), 3:1 ratio respectfully, and 2.5g/L Triton X-100 (Sigma, X100RS) at pH 6.5 in the presence of 0.1% BSA (Sigma, A8806). Rodent sample diluted in lysis buffer was mixed with equal volume of 4-Methylumbelliferyl-β-pyranoside (Sigma, M3633) prepared in 0.1M sodium acetate with 10g/L of BSA (Sigma, A8806) and 2.5g/L of Triton X-100 at pH4.5. Subsequently, the reaction mixture was incubated for 1 hour at 37°C and the reaction was terminated by adding stop solution (1M glycine-NaOH, pH12.5). GBA1 activity was determined by measuring cleavage of a synthetic substrate, 4-Methylumbelliferone, and release of a fluorophore. Fluorescence excitation at 365nm and emission at 445nm with a cutoff of 420nm were measured via Molecular Devices 96-well microtiter plate reader and SoftMax Pro version 5.4.4 software. GBA1 activity was reported as ng/ml/hr.

#### GCase activity assay

Equal amount of protein lysate from rodent and NHPs were used to determine GBA1 activity with glucosylceramidase activity assay kit (abcam, ab273339). GBA1 activity was determined by measuring cleavage of a synthetic substrate, 4-Methylumbelliferone, and release of a fluorophore. Fluorescence excitation at 360nm and emission at 445nm were measured via Molecular Devices SpectraMax M2e 96-well microtiter plate reader and SoftMax Pro version 7 software. GBA1 activity was reported as μU/mg (pmol/min/mg).

### In-house human Glucosylceramidase Enzyme-linked immunosorbent assay (ELISA)

96-well EIA/RIA Assay microplates (Corning, 9018) were coated with 2.5µg/mL GBA recombinant rabbit monoclonal Ab (Invitrogen, MA5-38382) in carbonate buffer (Invitrogen, CB01100) for overnight at 4°C. Wells were washed 3X in wash buffer (Invitrogen, WB01) and blocked in assay buffer (Invitrogen, DS98200) for overnight at 4°C. Standard curve was generated using recombinant human GBA protein (R&D Systems, 7410-GHB-020) for quantification of human GBA protein. Standards and samples were run in duplicate and incubated for 2 hours at room temperature, washed 3X in wash buffer, and then incubated with 2.5µg/mL recombinant biotin anti-GBA antibody (abcam, ab201496) for 2 hours at room temperature. Wells were washed 3X in wash buffer and incubated with 0.1µg/mL streptavidin HRP conjugated (Thermo Scientific, 21126) for 1 hour at room temperature. Subsequently, wells were washed 3X in wash buffer and incubated with TMB substrate (Invitrogen, EB02) for 30 minutes before adding stop solution (Invitrogen, SS04). Human GBA protein was quantified by measuring absorbance at 450nm with wavelength correction set to 540nm using Molecular Device SpectraMax and Softmax Pro version 7.1.2 software.

### Immunoblot preparation and analysis

Tissue homogenization, protein extraction, and protein quantification were previously described. For cell culture media, Pierce Protein Concentrators PES, 10K MWCO, 0.5mL (Thermo fisher, 88513) was used to concentrate 10-fold by following manufacturer’s protocol and the equal amount of protein was used for each sample. First, Protein samples were loaded on Criterion TGX stain-free precast gel (Biorad, 5678085) and transferred for 7min using Trans-Blot turbo midi 0.2µm PVDF transfer pack (Biorad, 1704157) and Trans-Blot Turbo Transfer System (Biorad, 1704150). After transfer, the membranes were blocked at room temperature for 2 hours with Licor Blocking Buffer (Licor, 927-60001) and followed by overnight blocking in Human Glucosylceramidase/GBA antibody (1:1000, R&D systems, MAB7410) and GAPDH antibody (1:5000, Cell signaling Technology, 2118) at 4°C. Then, the membranes were incubated in IR dye 800CW Donkey anti-mouse IgG (Licpr, 102673-332) and IR dye 680RD Donkey anti-rabbit IgG (Licor, 102673-414). Images are acquired using Licor Odyssey CLx imaging system (Licor, 9140).

### Lipid extraction

To quantify Glucosylsphingosine (GlcSph) and/or Glucosylceramide (GlcCer), 20μl of tissue homogenate (100mg/mL tissue weight) was aliquoted into a labeled 1.5ml Eppendorf tube followed by 180μl of internal standard solution (10ng/ml d5-GlcSph and 20ng/mL d35- C18GalCer in 30% methanol, 70% acetonitrile with 5mM ammonium acetate, and 1% acetic acid). The samples were vortexed for 10 minutes and sonicated for 10min. The tubes were sitting at 4°C for 10 minutes and centrifuged at 13,000 × g for 10 minutes. The supernatant (150uL) from each tube was transferred into a pre-labeled total recovery MS vial for MS analysis. Calibration curves for GlcSph and GlcCer were prepared in a pooled matrix, and concentrations were ranged from 0.03 to 1000 ng/ml.

### Lipid profiling by LC-MS/MS

The lipids were injected (5μl) into an LC/MS/MS system comprised of an Acquity UPLC (Waters, Milford, MA) and Sciex Triple Quad 5000 mass spectrometer (Sciex, Toronto, Canada). For GlcSph, the chromatographic separation (from GalSph) was achieved with a Waters Acquity BEH HILIC (2.1 x 100mm, 1.7mm particles, Part # 186003461) using mobile phases: (A) 96% ACN, 2% MeOH, 1% Acetic acid, 1% H2O, 5mM Ammonium acetate and (B) 98% MeOH, 1% Acetic acid, 1% H2O, 5mM Ammonium acetate. The column was maintained at 30°C. GlcSph was eluted with the following gradient: from 5% B to 50% B over 3 minutes, then the mobile phase composition was held constant for 0.5min followed by a rapid return (0.1 minute) to 5% B maintained for 1minute. All experiments were carried out at a flow rate of 0.5ml/min. Data were analyzed in Analyst (AB Sciex, Toronto, Canada). For GlcCer, the chromatographic separation (from GalCer) was achieved with a Waters Cortecs HILIC (2.1 x 100mm 2.7µm particles cat#186007427) using mobile phases: (A) 96% ACN, 2% MeOH, 1% Acetic acid, 1% H2O, 5mM Ammonium acetate and (B) 80% MeOH, 1% Acetic acid, 20% H2O, 5mM Ammonium acetate. The column was maintained at 20°C. GlcCer was eluted at an isobaric flow of 2% B for 4.5min. All experiments were carried out at a flow rate of 0.5ml/min. Data were analyzed in Analyst (AB Sciex, Toronto, Canada).

### Protein extraction for human GBA1 LC-MS

Lysis buffer containing 100mM TBS, pH 8.0, 100mM triethylammonium bicarbonate buffer, pH 8.0 (Sigma, 18597), 100mM Octyl Glucoside (Sigma, 10634425001), 2% (vol/vol) NP-40 (Thermo Scientific, 28324), 1mM PMSF (CST, 8553S), 1μM aprotinin (Sigma, A3428), 1X phosphatase inhibitor (CST, 5870S) was added to NHP tissues as 5 times the dry weight of tissues. Tissues were homogenized in 1.4mm ceramic bead tubes (Fisher Scientific, 15-340- 153) using OMNI Ruptor 12 (OMNI International). Following homogenization and centrifugation at 18000 g at 4°C, clear lysates were collected and subjected to BCA (Thermo Scientific, 23225) to measure protein concentration.

### LC-MS analysis of hGBA1 protein

Tissue homogenates from human or non-human primates (NHP) were prepared for proteomic analysis. This involved immunoprecipitation using protein A/G beads and four antibodies against GBA (two monoclonal: Novus NBP2-45829 and Abnova H00002629-M01; and two polyclonal: Sigma G4046 and Invitrogen PA5-67940). The samples underwent stepwise reduction, alkylation, and overnight digestion with Trypsin/Lys C and ArgC proteases. High-throughput sample preparation was facilitated by the MINI 96 digital pipettor (Integra Biosciences).

The digested peptides, spiked with synthetic heavy-labeled peptides (C-terminal R/K (^13^C^15^N)), were clarified using EvoTip trap columns. These were connected online with an Evo 8-cm analytical column and separated on an 11.5-minute LC gradient (processing 100 samples per day) using the Evosep One nanoLC system. Absolute quantification of GBA proteins was attempted using an in-house multiplex LC-MS assay, which employed high-field asymmetric waveform ion mobility spectrometry (FAIMS) for gas-phase ion separation in high-resolution mode. FAIMS-separated ions were analyzed via parallel reaction monitoring (PRM), with compensation voltages (CV) for target peptides optimized by adjusting DC voltages from -50CV to -30CV to identify the best-performing CV values.

Human and NHP brain homogenates were processed and analyzed similarly. MS1 data for all samples were acquired at high resolution (240k), and tMS2 data at 30k resolution. The accuracy and precision of FAIMS-PRM assays were evaluated using a 9-point calibration curve, with a linear range of 20 ng to 11 μg/mL for the human-specific analyte PVSLLASPWTSPTWLK, and 40 ng to 11 μg/mL for the common human and NHP analyte FWEQSVR. Quality controls (QCs) at low, medium, and high levels contained surrogate peptides spiked into a pooled matrix. The ratio of endogenous to surrogate peptides from FAIMS-PRM data was obtained using Skyline software, and GBA expression values were calculated using a single-point calibration method from the equation as follows:

R = C_GBA_ (fmol/μL or nM) = [{(R x S_O_)/V_d0_} x T_Vd_] ÷ B_Vd_

= L/H; L= endogenous and H= surrogate (heavy)
S_O_ = spike (H) on-column (fmol)
V_d0_ = digests volume (on-column)
T_Vd_ (μL) = total volume (μL) of starting matterials
B_Vd_ (μL) = Total volume (μL) of brain homogenate (undigested) provided

### Chromogenic *In situ* Hybridization (ISH)

RNAScope In Situ Assay for WPRE was performed using RNAScope 2.0 Brown detection kit (ACD, 320497). Rodent and NHP brain FFPE slides were pre-treated in EDTA Buffer (pH 9.0) at 90°C followed by protease treatment at 40°C. The RNAScope WPRE ISH (ACD, 410058) assay was carried out in 40°C oven for hybridization and amplification from Amp 1 to Amp 4, followed by Amp 5 and Amp 6 at room temperature. The WPRE ISH signal was developed using DAB producing brown color.

### Chromogenic Immunohistochemistry (IHC)

Rodent and NHP brain FFPE slides were treated in EDTA solution (pH 9.0) for antigen retrieval and subsequently blocked with 3% hydrogen peroxide, followed with 5% horse serum. The slides were then incubated with human GBA1 antibodies (1:100, abcam, ab125065 and 1:400, Novus, NBP2-45829) for rodent and NHP, respectively. The color development for GBA1 signal was achieved by DAB resulting brown color staining and the slides were counter stained with hematoxylin for nuclei staining.

### Immunofluorescent RNA/Protein integrated co-detection assay

Co-detection of WPRE mRNA with payload human GBA protein and different cell type markers from NHP brain FFPE slides were performed using Leica BOND RX automated stainer (Leica Biosystems). Multiplexing protocol was programmed for streamline In Situ Hybridization and a single continuous IHC staining for co-detection of RNA and protein markers. RNAscope 2.5 LS multiplex fluorescent kit (ACD, 322800) and Bond polymer refine detection kit (Leica Biosystems, DS9800) were used in this new programmed protocol. WPRE probe (ACD, 410058) was used in In Situ Hybridization step followed by detection and visualization using TSA Vivid Fluorophore 570 (1:1000, biotechne, 323272). In sequential IHC steps, GBA1 antibody (1:6000, Novus, NBP2-45829) and cell type markers (1:500, ab177487, NeuN from abcam, 1:500, 42397, S100b from CST, 1:500, 17198, Iba-1 from CST, and 1:500, ab109186, Olig2 from abcam) were used for detection of human GBA and cell type markers. DAPI signal was used as nuclear counterstaining.

### NHP sample collection

At necropsy, animals were perfused with chilled PBS pH 7.4, and their brains were cut in a brain matrix into 4 mm coronal slices and hemisected. The left hemispheres were frozen on dry ice for biochemical analysis, while the right hemispheres were fixed in 10% neutral buffered formalin for 36-48 hours at room temperature before embedding in paraffin blocks for histopathology and immunohistochemistry analysis. The spinal cord and DRGs from 4 levels (cervical, upper thoracic, lower thoracic and lumbar), as well as peripheral tissues were collected as frozen and fixed samples. Tissue punches were collected using 3 mm biopsy punches from indicated brain regions on the frozen brain slices, as well as from 4 levels of spinal cords, DRGs and peripheral tissues. Paraffin tissue sections were stained for hematoxylin and eosin and submitted for histopathology analysis.

### NHP histopathology

Pathology evaluations were conducted by a board-certified veterinary pathologist at HistoWiz (Long Island City, NY) on hematoxylin and eosin-stained slides. For each animal, eight key brain regions (cerebral cortex, basal ganglia, thalamus, hippocampus, midbrain/pons, cerebellum, medulla and major white matter tracts), as well as spinal cord and DRGs from cervical, upper thoracic, lower thoracic, and lumbar levels were examined for axon/neuron degeneration and/or inflammation/cell infiltration. Microscopic findings were graded as 0 for absence of lesion, 1 for minimal, 2 for mild, 3 for moderate, 4 for marked, and 5 for severe.

### Vector genome assessment

#### Rodent or NHP tissue homogenates

gDNA was isolated using QIAamp 96 DNA QIAcube HT kit (QIAGEN, 51331) according to manufacturer’s protocol. Briefly, tissue homogenates were treated with proteinase K at 56°C for overnight and transferred to S block (QIAGEN, 19585). The samples were placed into QIAGEN QIAcube HT instrument and gDNA isolation was performed by following the steps from QIAcube HT Prep Manage Software.

#### Rodent bone marrow

After gDNA isolation, the concentration was measured with NANODROP 8000 (Thermo Fisher Scientific) and vector genome was determined by dPCR with QIAGEN QIAcuty instrument. The gDNA samples were analyzed using primer-probe combination specific to the bGH, transgene, and housekeeping gene sequences (Integrated DNA Technology) to determine the vector copy number.

### Transcript assessment

RNA was isolated from rodent or NHP tissue homogenates using RNeasy 96 QIAcube HT kit (QIAGEN, 74171) according to manufacturer’s protocol. Briefly, tissue homogenates were mixed with QIAzol lysis reagent (QIAGEN, 79306) and followed by chloroform (Fisher Scientific, C298- 1). The mixture was centrifuged at 4°C and the aqueous phase was transferred to S block (QIAGEN, 19585) for further RNA isolation. The samples were placed into QIAcube HT instrument and RNA isolation was performed by following the steps from QIAcube HT Prep Manage Software with in-column DNase (QIAGEN, 79256) treatment. After RNA isolation, the concentration was measured with NANODROP 8000 (Thermo Fisher Scientific). RT-dPCR was carried out by producing cDNA via QIAcuity OneStep advanced probe kit (QIAGEN, 250132) and using primer-probe combination specific to the transgene, GFP, and housekeeping gene sequences (Integrated DNA Technology) to determine the total transcripts per RNA input.

### NIH NeuroBioBank

Post-mortem human tissues were procured from the NIH NeuroBioBank at the University of Maryland Brain and Tissue Bank and Sepulveda Research Corporation. Selected human donors were 55-70 years old with no pathological CNS clinical records or diagnosed neurodegenerative disorders. Twelve distinct regions were selected to represent different functional brain regions.

### Statistics

All statistical analyses were performed with GraphPad Prism 9 software (GraphPad, San Diego, CA, USA) using either one-way or two-way ANOVA with Tukey’s multiple comparisons test or with Student’s *t* test depending on the data set.

## Acknowledgements

We thank the following people for their support in this project: AAV manufacturing done by Shelley Nass (Sanofi) and Khushdeep Mangat (Sanofi), Hyejung, Catie Viel, Amy Richards, Kasey Jackson (Sanofi) for rodent CBE studies, statistics review by Weilian Qiu (Sanofi), project administration support by Kristin Radzwill (Sanofi), portfolio supervision by Evis Havari (Sanofi), IP support by Alejandro Martinez (Sanofi). Review and writing support by Michael Fleming (Sanofi) and Edith Pfister (Sanofi).

## Author contributions

Conceptualization: SA, SR, CM, PS

Experiments, methods and data generation: SA, JR, BC, CVM, LB, EW, JB, LG, MH, DG, WM, AB, LC, JM, CK

Data review: SA, CK, PS, MG, BE, SR Writing: SA, BE, SR

## Data and materials availability

All data are available upon reasonable request from.

