## Supplemental Figures and Tables for "AAV gene therapy for GBA-PD and Gaucher Disease"

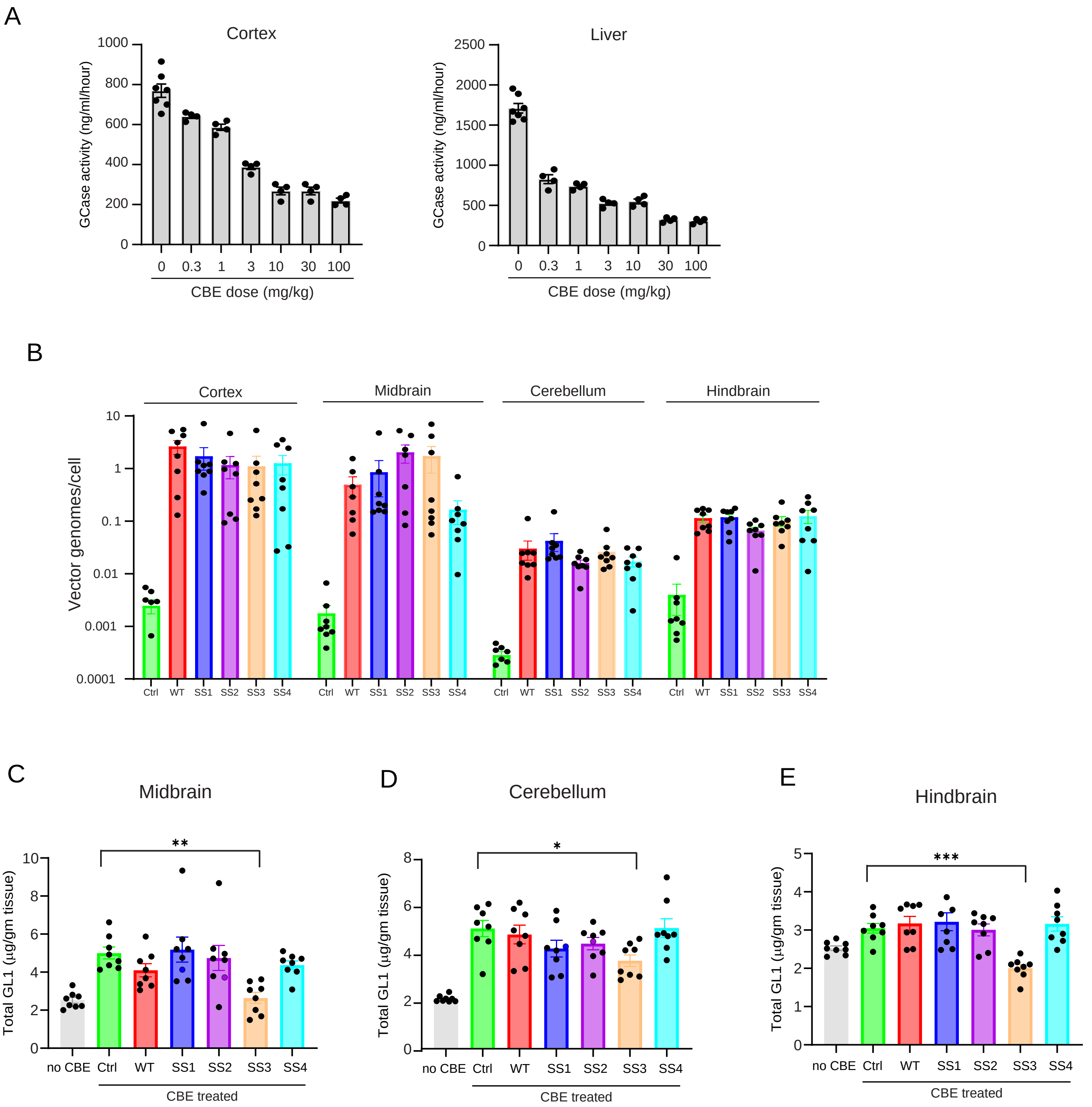
A
Cortex
GCase activity (ng/ml/hour)
1000
800
600
400
200
0
0 0.3 1 3 10 30 100
CBE dose (mg/kg)
Liver
GCase activity (ng/ml/hour)
2500
2000
1500
1000
500
0
0 0.3 1 3 10 30 100
CBE dose (mg/kg)
B
Cortex
Midbrain
Cerebellum
Hindbrain
Vector genomes/cell
10
1
0.1
0.01
0.001
0.0001
Ctrl WT SS1 SS2 SS3 SS4
C
Midbrain
Total GL1 (µg/gm tissue)
**
no CBE Ctrl WT SS1 SS2 SS3 SS4
CBE treated
D
Cerebellum
*
E
Hindbrain
***

**Figure S1. CBE-induced Lipid flux and efficient lipid substrate clearance by SS3-GBA1 in WT mice**

(A) GCase enzyme activity measured in cortex and liver from CBE-induced C57 WT/BL6 mouse. N=7 animals for CBE 0 mg/kg and N=4 per all the other groups. Data are Mean ± SEM

(B) AAV vector genome copies measured in four different brain regions by bGH dPCR and normalized to the *Rab1a* gene intron to obtain VG copies per cell.

(C - E) Measurement total GL-1 lipid substrate levels by LC-MS in midbrain (C), cerebellum (D), and hindbrain (E). No CBE group in the graphs shown is control mice that did not receive any AAV vector or CBE injection. 8 mice per group. ***$p<0.001$, **$p<0.01$, *$p<0.05$; One way-ANOVA with Tukey's multiple comparisons test with all groups compared to CBE-treated vehicle injected group.

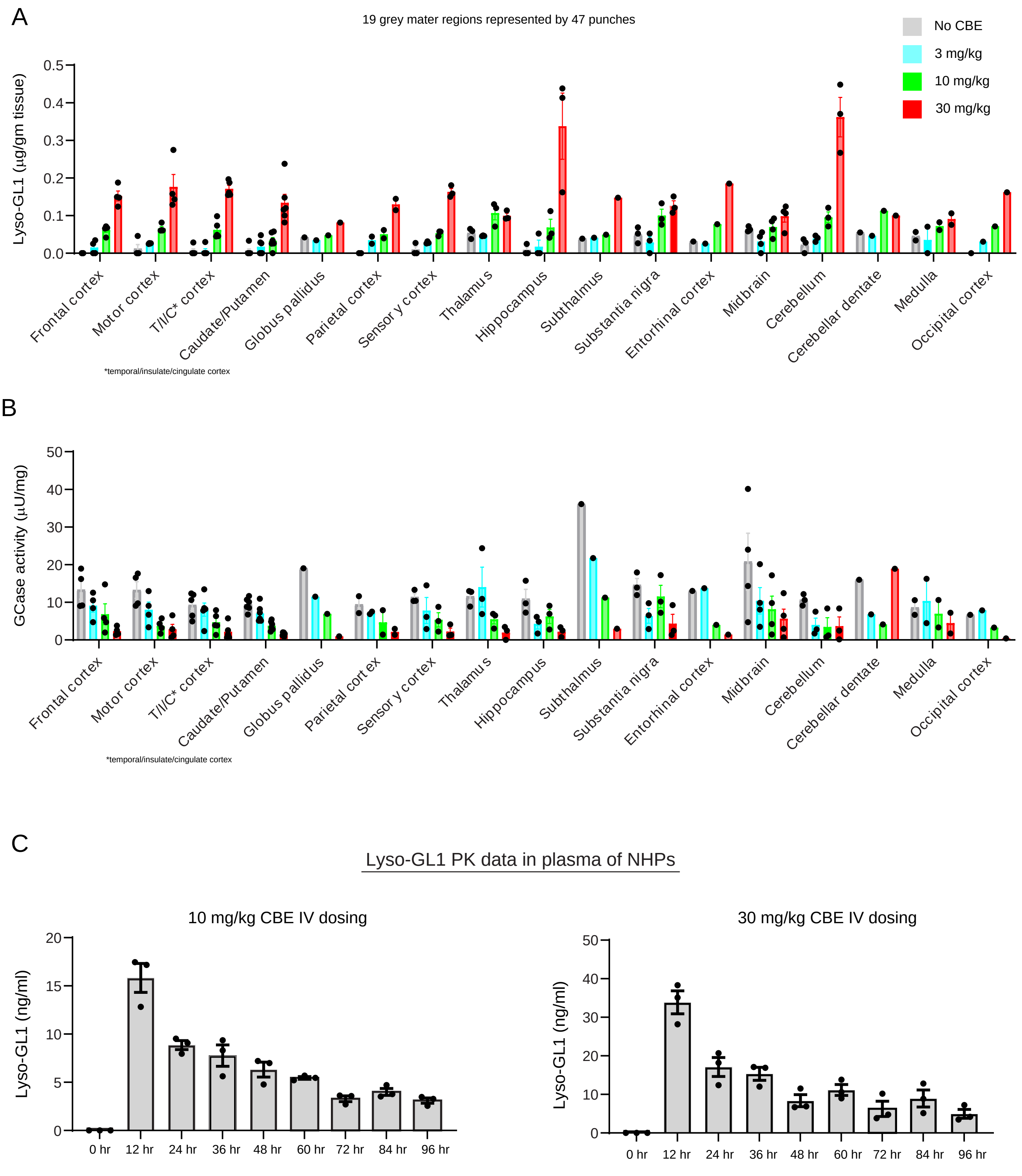
A
19 grey mater regions represented by 47 punches
No CBE
3 mg/kg
10 mg/kg
30 mg/kg
Lyso-GL1 (µg/gm tissue)
0.0
0.1
0.2
0.3
0.4
0.5
Frontal cortex
Motor cortex
T/I/C* cortex
Caudate/Putamen
Globus pallidus
Parietal cortex
Sensor y cortex
Thalamus
Hippocampus
Subthalmus
Substantia nigra
Entorhinal cortex
Midbrain
Cerebellum
Cerebellar dentate
Medulla
Occipital cortex
*temporal/insulate/cingulate cortex
B
GCase activity (µU/mg)
0
10
20
30
40
50
Frontal cortex
Motor cortex
T/I/C* cortex
Caudate/Putamen
Globus pallidus
Parietal cort ex
Sensor y cortex
Thalamus
Hippocampus
Subthalmus
Substantia nigra
Entorhinal cortex
Midbrain
Cerebellum
Cerebellar dentate
Medulla
Occipital cortex
*temporal/insulate/cingulate cortex
C
Lyso-GL1 PK data in plasma of NHPs
10 mg/kg CBE IV dosing
Lyso-GL1 (ng/ml)
0
5
10
15
20
0 hr
12 hr
24 hr
36 hr
48 hr
60 hr
72 hr
84 hr
96 hr
30 mg/kg CBE IV dosing
Lyso-GL1 (ng/ml)
0
10
20
30
40
50
0 hr
12 hr
24 hr
36 hr
48 hr
60 hr
72 hr
84 hr
96 hr

**Figure S2. Development of CBE-induced lipid flux model in NHPs**

(A and B) Total of 47 brain biopsy punches surveyed at the individual brain region level encompassing 19 grey matter regions. Lyso-GL1 level (A) and GCase enzyme activity (B) were measured. Data are Mean ± SEM. T/I/C cortex: temporal/insulate/cingulate cortex

(C) Pharmacokinetic analysis of Lyso-GL1 in plasma of NHP with 10 mg/kg CBE and 30 mg/kg CBE by IV dosing. Data are Mean ± SEM. 3 animal at each time point.

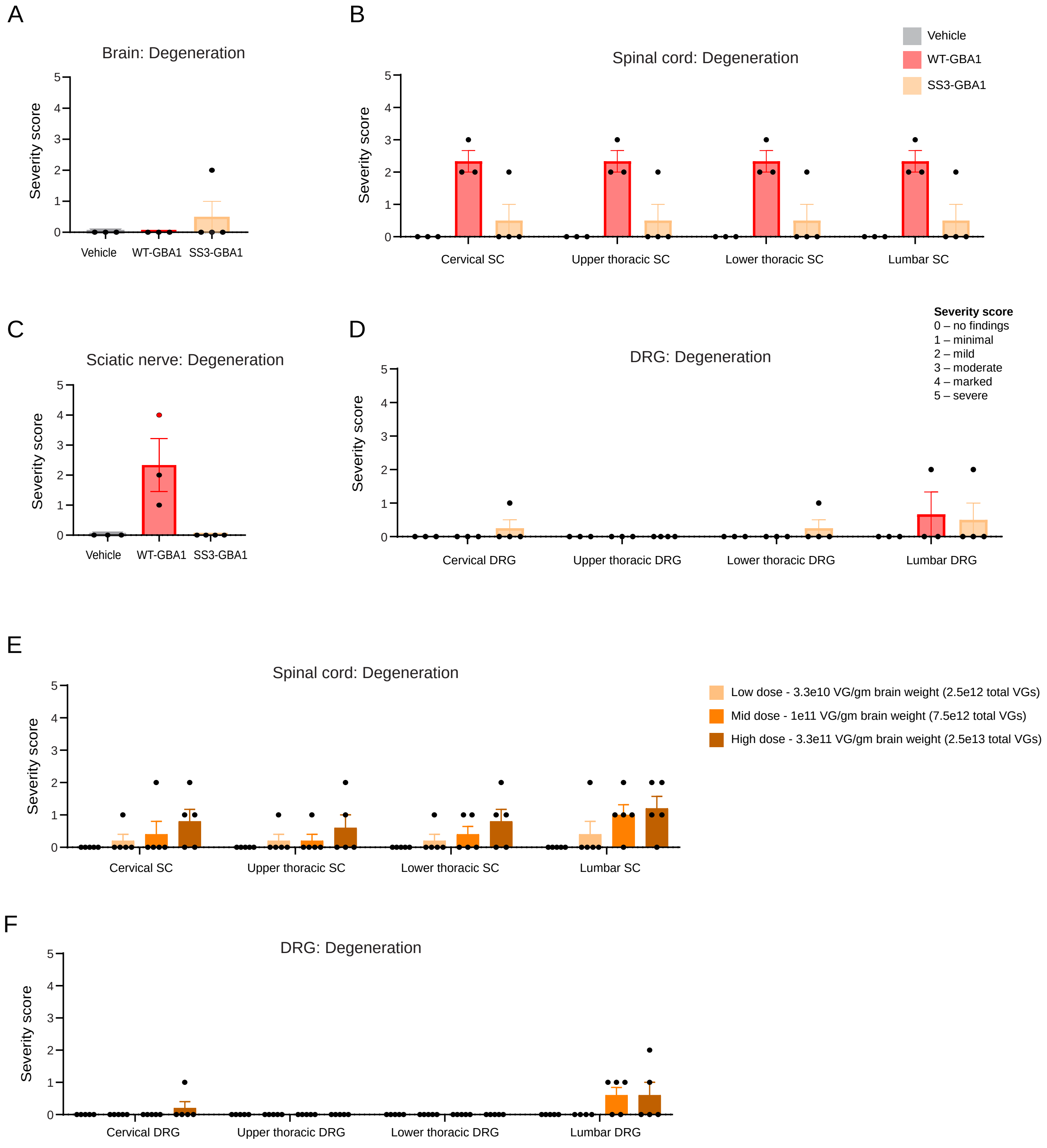
A
Brain: Degeneration
Severity score
0
1
2
3
4
5
Vehicle
WT-GBA1
SS3-GBA1
B
Spinal cord: Degeneration
Vehicle
WT-GBA1
SS3-GBA1
Severity score
Cervical SC
Upper thoracic SC
Lower thoracic SC
Lumbar SC
C
Sciatic nerve: Degeneration
Severity score
Vehicle
WT-GBA1
SS3-GBA1
D
DRG: Degeneration
Severity score
0 – no findings
1 – minimal
2 – mild
3 – moderate
4 – marked
5 – severe
Severity score
Cervical DRG
Upper thoracic DRG
Lower thoracic DRG
Lumbar DRG
E
Spinal cord: Degeneration
Low dose - 3.3e10 VG/gm brain weight (2.5e12 total VGs)
Mid dose - 1e11 VG/gm brain weight (7.5e12 total VGs)
High dose - 3.3e11 VG/gm brain weight (2.5e13 total VGs)
Severity score
Cervical SC
Upper thoracic SC
Lower thoracic SC
Lumbar SC
F
DRG: Degeneration
Severity score
Cervical DRG
Upper thoracic DRG
Lower thoracic DRG
Lumbar DRG

**Figure S3. Safety profile of AAV GMU01-SS3-GBA1 in NHP studies**

Histopathology findings reported with severity scores across central nervous system tissues by board certified clinician.

(A – D) SS3-GBA1 exhibited better safety profile than WT-GBA1. Scores reported for brain (A), spinal cord (B), sciatic nerve (C), and DRGs (D). Data are Mean ± SEM. Each dot is score for individual NHP. 3 - 4 NHPs per group.

(E and F) Scores reported for spinal cord (E) and DRGs (F) from NHP dose-range finding study with minimal to mild histopath finding for degeneration. Data are Mean ± SEM. Each dot is score for individual NHP. 5 NHPs per dosed group.

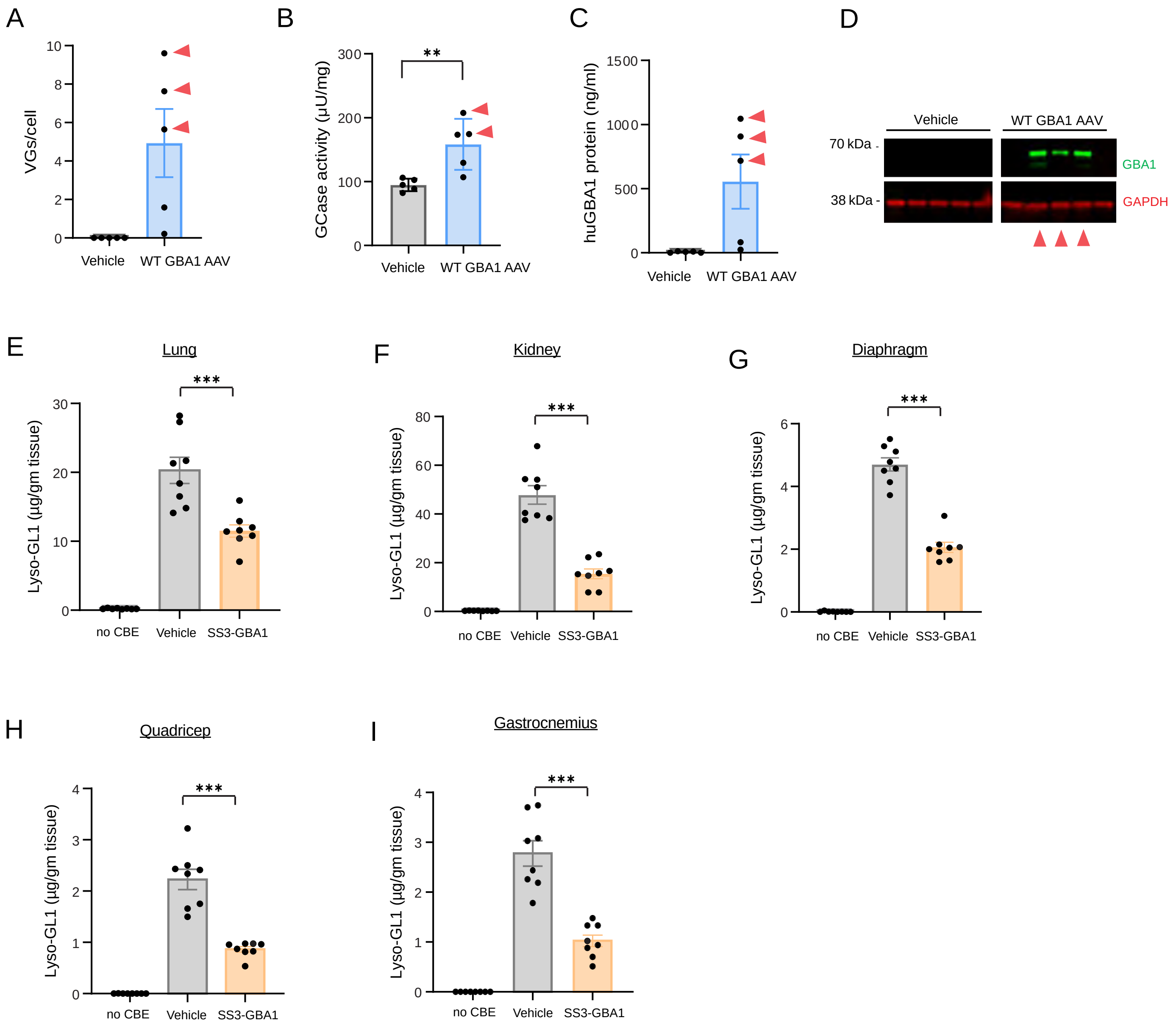

A
VGs/cell
Vehicle
WT GBA1 AAV
B
GCase activity (µU/mg)
**
Vehicle
WT GBA1 AAV
C
huGBA1 protein (ng/ml)
Vehicle
WT GBA1 AAV
D
Vehicle
WT GBA1 AAV
70 kDa -
38 kDa -
GBA1
GAPDH
E
Lung
Lyso-GL1 (µg/gm tissue)
***
no CBE
Vehicle
SS3-GBA1
F
Kidney
Lyso-GL1 (µg/gm tissue)
***
no CBE
Vehicle
SS3-GBA1
G
Diaphragm
Lyso-GL1 (µg/gm tissue)
***
no CBE
Vehicle
SS3-GBA1
H
Quadricep
Lyso-GL1 (µg/gm tissue)
***
no CBE
Vehicle
SS3-GBA1
I
Gastrocnemius
Lyso-GL1 (µg/gm tissue)
***
no CBE
Vehicle
SS3-GBA1

**Figure S4. Validation of in-house human GBA1 ELISA and lipidomics in tissues from intravenous dosing of SS3-GBA1 in mice**

In-house sandwich ELISA was developed to measure human GBA1 protein levels in bulk tissues. For validation, brain tissue samples were used from a study where vehicle or AAV GMU01-WT-GBA1 were administered by bilateral ICV (A/P: -0.3, M/L: +/-1.0, D/V: -2.0) to 4-month-old CT57 WT/BL6 mice at 1.0e11 vector genomes per animal and 5µl per hemisphere. 5 animals per group.

(A – D) 4 weeks post AAV injection, brain tissue samples were analyzed for vector genome (A), GCase enzyme activity (B), and human GBA1 protein level by ELISA (C) and western blotting (D). Data are Mean ± SEM. 5 animals per group. 3 mice with higher amount of VGs/cell track similarly across GCase activity and protein quantification assays.

(E – I) Lyso-GL1 lipid clearance in SS3-GBA1 injected mice compared to vehicle injected in lung (E), kidney (F), diaphragm (G), quadricep (H) and gastrocnemius (I). 8 animals per group, No CBE group in the graphs shown is control mice that did not receive any AAV vector or CBE injection. Data are Mean ± SEM. ***$p<0.0001$; One way-ANOVA with Dunnett's multiple comparisons test (Lyso-GL1). All groups compared to vehicle group.

**Table S1**

| Transgene name | Likelihood of signal peptide | Cleavage probability | DeepLoc 1.0 prediction | Lysosome likelihood | Extracellular likelihood |
|---|---|---|---|---|---|
| WT-GBA1 | 0.9979 | 0.788 | Lysosome, soluble | 0.8688 | 0.0898 |
| SS1-GBA1 | 0.9997 | 0.977 | Extracellular, soluble | 0.3029 | 0.6832 |
| SS2-GBA1 | 0.9997 | 0.974 | Extracellular, soluble | 0.3563 | 0.6069 |
| SS3-GBA1 | 0.9998 | 0.98 | Extracellular, soluble | 0.3537 | 0.6212 |
| SS4-GBA1 | 0.9994 | 0.9664 | Extracellular, soluble | 0.3381 | 0.6458 |

**Table S2**

| Antibodies | Vendor | Catalog number | Application |
|---|---|---|---|
| Rabbit monoclonal anti-GBA [EPR5142] | abcam | ab125065 | WB, ICC IF, IHC |
| GBA Recombinant Rabbit Monoclonal Antibody (4H4) | Invitrogen | MA5-38382 | ELISA |
| Biotin Anti-GBA antibody [EPR5142] | abcam | ab201496 | ELISA |
| Mouse monoclonal Human Glucosylceramidase/GBA Antibody | R&D systems | MAB7410 | WB |
| Rabbit monocolonal anti-GAPDH, clone 14C10 | Cell Signaling Technology | 2118 | WB |
| Rabbit monoclonal anti-LAMP1A | Cell Signaling Technology | 9091 | ICC IF |
| Mouse monoclonal anti-GBA [OTI4G4] | Novus Biologicals | NBP2-45829 | ISH/IHC IF Multiplexing, IP-MS |
| GBA monoclonal antibody (M01), clone 2E2 | Abnova | H00002629-M01 | IP-MS |
| Anti-Glucocerebrosidase antibody produced in rabbit | Sigma | G4046 | IP-MS |
| GBA Polyclonal Antibody | Invitrogen | PA5-67940 | IP-MS |
| Rabbit monoclonal anti-NeuN [EPR12763] | abcam | ab177487 | ISH/IHC IF Multiplexing |
| Rabbit monoclonal anti-S100B [E7C3A] | Cell Signaling Technology | 42397 | ISH/IHC IF Multiplexing |
| Rabbit monoclonal anti-Iba-1 [E4O4W] | Cell Signaling Technology | 17198S | ISH/IHC IF Multiplexing |
| Rabbit monoclonal anti-Olig2 [EPR2673] | abcam | ab109186 | ISH/IHC IF Multiplexing |
